# Induced Degradation of Lineage-specific Oncoproteins Drives the Selective PARP1 Inhibitor Toxicity in Small Cell Lung Cancer

**DOI:** 10.1101/2022.11.02.514072

**Authors:** Chiho Kim, Xu-Dong Wang, Shuai Wang, Peng Li, Zhenzhen Zi, Qing Ding, Seoyeon Jang, Jiwoong Kim, Yikai Luo, Kenneth E. Huffman, Ling Cai, Han Liang, John D. Minna, Yonghao Yu

**Author notes:** Department of Molecular Pharmacology and Therapeutics, Columbia University Irving Medical Center, New York, NY 10032, USA. These authors contributed equally (Chiho Kim, Xu-Dong Wang). Correspondence (Y.Y.).

## Abstract

A subset of small cell lung cancer (SCLC) shows clinically relevant response to PARP1 inhibitors (PARPi). However, *BRCA1/2* mutations are not commonly found in SCLC, and the underlying mechanism(s) of PARPi sensitivity in SCLC is poorly understood. We performed quantitative proteomic analyses and identified proteomic changes that signify PARPi responses in a large panel of molecularly annotated patient-derived SCLC lines. We found that the toxicity of PARPi in SCLC could be explained, at least in part, by the PARPi-induced degradation of key lineage-specific oncoproteins including ASCL1, NEUROD1, POU2F3, KDM4A, and KDM5B. Importantly, the degradation of these SCLC lineage-specific oncoproteins could also be induced by commonly used chemotherapeutic agents. Biochemical experiments showed that PARPi-induced activation of E3 ligases (e.g., HUWE1 and RNF8) mediated the ubiquitin-proteasome system (UPS)-dependent degradation of these oncoproteins. Interestingly, although PARPi resulted in a general DNA damage response in SCLC cells, this signal is sensed by different SCLC cell lines to generate a cell-specific response. The dissection of the cell-specific oncoprotein degradation response led to the identification of potentially predictive biomarkers for PARPi in SCLC. The combination of PARPi and agents targeting these pathways led to dramatically improved cytotoxicity in SCLC. PARPi-induced degradation of lineage-specific oncoproteins therefore represents a novel mechanism to explain the efficacy of PARPi in tumors without *BRCA1/2* mutations.

**Highlights:**

1. Quantitative mass spectrometric analysis identifies proteomic changes associated with PARPi treatment in a large panel of SCLC cell lines.
2. PARPi leads to the degradation of lineage-specific oncoproteins (e.g., ASCL1 and KDM4A) via the DNA damage responsive E3 ubiquitin ligases (e.g., HUWE1 and RNF8).
3. A combination of PARPi and agents targeting the lineage-specific oncoproteins offers a more complete and durable therapeutic response in SCLC, compared to PARPi alone.
4. Expression of lineage-specific oncoproteins and the associated ubiquitination machinery are predictive biomarkers for PARPi-induced cytotoxicity in SCLC.

## Introduction

Poly(ADP-ribose) polymerase 1 (hereafter referred to as PARP1) is one of the PARP enzymes that is critically involved in DNA damage response (DDR). Upon the binding to nicked DNA, PARP1 becomes activated to synthesize many Poly-ADP-ribosylated (PARylated) proteins, including itself. These protein-linked PAR polymers serve as a scaffold to recruit an array of DNA repair enzymes to resolve the DNA breaks^1, 2^. Besides the blockade of PAR synthesis and PAR-mediated DNA damage response, recent studies show that PARP1 inhibitor (PARPi) also causes PARP1 trapping^3–5^. PARP1 auto-PARylation triggers the release of PARP1 from the DNA lesions, owing to the steric hindrance and charge repulsion introduced by the PAR polymers. Thus, PARPi blocks this key step in PARP1-dependent DDR, causing the formation of toxic PARP1-PARPi complexes at the DNA damage site. These complexes interfere with DNA replication, resulting in the stalling and collapse of the replication fork, the conversion of single-strand breaks (SSB) to double-strand breaks (DSB), and eventually, cell death. Recent studies have also shown that PARP1 trapping is a key mediator of the immunomodulatory roles of PARPi^5, 6^.

Cancer cells with defects in the homologous recombination repair (HRR) pathways, such as *BRCA1/2*-mutated cells, are particularly sensitive to PARPi. They are more susceptible to PARP1 trapping, because of the lack of the HRR mechanism to remove the deleterious effects of PARP1 trapping (known as the synthetic lethality)^7^. Indeed, dramatic responses to single-agent PARPi treatment were observed in *BRCA1/2*-mutated breast and ovarian cancer patients, and these results led to the revolutionized management of these patients, with the recent FDA approval of multiple PARPi for these indications^7, 8^.

Besides these very exciting results in *BRCA1/2*-mutated breast and ovarian cancers, PARPi is being evaluated, either as a single agent or in combination with chemotherapy and radiotherapy approaches, to treat a diverse array of other solid tumors. The results from these studies showed that several of these tumor types (e.g., small cell lung cancer, SCLC) displayed increased sensitivity to PARPi. SCLC is a neuroendocrine (NE) lung carcinoma (~15% of all lung cancer cases) and is one of the most aggressive human malignancies (5-year survival rate of ~6%)^9^. Compared to other subtypes of lung cancer (e.g., adenocarcinoma and squamous cell carcinoma), SCLC has unique biology and genetic alterations, including the frequent deletion or mutation of the retinoblastoma (Rb) protein and p53 (TP53) protein. Despite the intensive research within the last several decades, the standard of care for advanced SCLC relies on chemotherapy, i.e., etoposide or topotecan plus a platinum-based drug such as cisplatin or carboplatin^10, 11^. Although this regimen often leads to initial tumor regression (in 60 to 80% of the cases), recurrence is almost invariable, and patients usually are left with very limited options of further systemic therapy^12, 13^. Therapeutic strategies that achieve a more complete and durable response for SCLC patients are therefore a significant unmet need. Despite the promising effects of PARPi in SCLC, mutations of *BRCA1/2* and other HRR genes are not commonly found in this tumor type (e.g., less than 3% of the SCLC cases contain *BRCA1/2* mutations)^14, 15^. These results raise the hypothesis that *BRCA1/2* mutations might not be the only determinant for PARPi sensitivity, at least in SCLC. The exact molecular underpinnings of PARPi toxicity in these tumors are poorly understood. The lack of mechanism-based, predictive biomarkers poses a significant challenge, preventing the broader application of PARPi in SCLC, as well as many other human malignancies with proficient HRR pathways.

A powerful approach to understanding the mechanism of action (MoA) for small molecule drugs is to perform systematic perturbation experiments and then monitor the adaptive responses^16–18^. Here, we performed isobaric labeling-based quantitative mass spectrometry experiments to comprehensively characterize how the SCLC proteome is remodeled upon the treatment of PARPi. This system-wide proteomic approach generated a large dataset of quantitative protein profiling to PARPi in SCLC. Among the various down-regulated proteins, we identified multiple lineage-specific oncoproteins, including ASCL1, NEUROD1, and POU2F3, and members of the histone demethylases, including KDM4A and KDM5B. We further uncovered the UPS (ubiquitin-proteasome system)-dependent mechanisms that mediate the PARPi-induced degradation of these lineage-specific oncoproteins. Furthermore, the combination of PARPi and agents targeting these pathways led to dramatically improved cytotoxicity in SCLC. Even though PARPi treatment elicits a general DNA damage signal, various SCLC cells responded differently to this DDR signal, leading to the unequal degradation of the lineage-specific oncoproteins, and hence cell death. The cell-specific oncoprotein degradation response and the associated heterogenous UPS response led to the identification of potentially predictive biomarkers for PARPi in SCLC. We expect that the dataset will serve as an invaluable resource to provide the foundation and biomarker for future hypothesis-driven research that helps delineate the molecular mechanisms of the cytotoxicity of PARPi beyond tumors with *BRCA1/2* mutations.

## Results

### Identification of a PARPi-induced Protein Degradation Signature (PiPS) in SCLC

*BRCA1/2* mutations are rarely found in SCLC, and the selective toxicity of PARPi in SCLC is poorly understood. Talazoparib is a highly potent PARPi that blocked the formation of PARylation in NCI-H2081 (H2081) cells (a representative SCLC cell line) (Supplementary Fig. 1A). Besides its catalytic inhibition activity, talazoparib treatment also led to profound PARP1 trapping, γH2AX formation (a marker of DNA double-strand breaks) as well as PARP1 cleavage (a marker of apoptotic cell death) (Supplementary Fig. 1B and C).

SCLC can be divided into different subtypes based on the expression of certain lineage-specific transcription factors, including ASCL1, NEUROD1, and POU2F3. Among them, the basic helix–loop–helix (bHLH) transcription factor ASCL1 is considered as a master regulator for a majority of these SCLC with strong neuroendocrine features. In addition, two related transcription factors, NEUROD1 and POU2F3, characterize a smaller subset with intermediate and low neuroendocrine features, respectively^19^. These transcription factors are highly expressed and are required for the establishment of the lineage of pulmonary neuroendocrine cells and for the continued survival of SCLC^20–23^.

In order to systemically characterize the proteomic response of SCLC to PARPi, we assembled a panel of 24 patient-derived SCLC cell lines that represented the major SCLC subtypes, including ASCL1^high^, NEUROD1^high^, POU2F3^high^ and TF^low^ (the TF^low^ subtype refers to the one that expresses low levels of ASCL1, NEUROD1 and POU2F3) (Supplementary Table S1). We first determined the sensitivity of these SCLC cell lines against talazoparib by measuring their IC_50_ (the half-maximal inhibitory concentration).

Talazoparib displayed differential toxicity in these SCLC cell lines, and we further classified these SCLC cell lines into the talazoparib-sensitive cells (IC_50_ < 1 μM, i.e., H2081, H69, H1048, H209, H1876, DMS-79, H2107, H524, H1092, H2171, H128, and H446) and talazoparib-resistant cells (IC_50_ > 1 μM, i.e., H1436, H82, H889, H378, H196, SW1271, H1836, H1341, H2029, H841, SHP-77, and H1963) (Fig. 1A; Supplementary Fig. 1D and E; Supplementary Table S1).

**Fig. 1.**
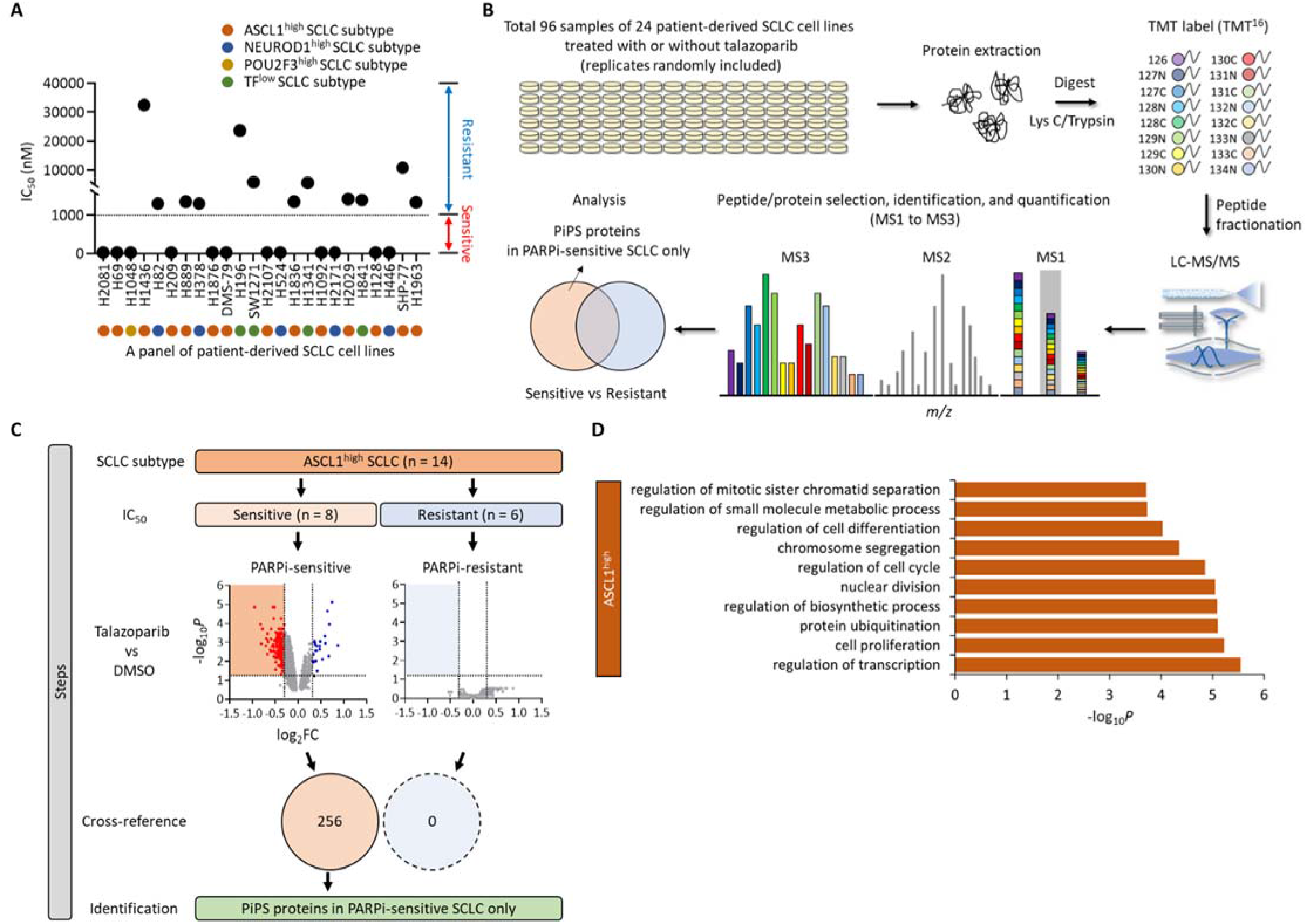
Identification of a PARPi-induced Protein Degradation Signature (PiPS) in SCLC. (A) The value of IC_50_ in a total of 24 SCLC cell lines treated with talazoparib (n = 3). SCLC cell lines are indicated as ASCL1^high^, NEUROD1^high^, POU2F3^high^, or TF^low^ subtype. The sensitivity is defined as: Sensitive (Red), IC_50_ < 1 μM; Resistant (Blue), IC_50_ > 1 μM. TF^low^, low expression of all three transcription factors. See also Supplementary Fig. 1A and B. (B) Workflow of high-throughput multiplexed quantitative proteome mapping in a total of 96 proteome samples including a total of 24 SCLC lines treated with or without talazoparib (1 μM for 48 hrs) in a total of 6 TMT experiments. See also Supplementary Table S2. (C) Steps of the identification of PiPS proteins only identified in PARPi-sensitive ASCL1^high^ SCLC subtype. Volcano plots show differentially expressed proteins in talazoparib treatment in each PARPi-sensitive and - resistant ASCL1^high^ SCLC subtype (Log2FC < −0.3, *P* < 0.05). Venn diagram shows the number of identified PiPS proteins. See also Supplementary Table S3. (D) Gene Ontology (GO) analyses of PiPS proteins identified in PARPi-sensitive ASCL1^high^ SCLC subtype. Biological processes were analyzed using the ToppGene database (https://toppgene.cchmc.org/).

We next performed quantitative mass spectrometric analyses to systemically characterize how talazoparib treatment perturbed the proteome homeostasis in each of the aforementioned SCLC cell lines (Fig. 1B). We prepared a total of 96 proteome samples (SCLC cell lines treated with either DMSO or talazoparib), and these samples were divided into 6 TMT-16plex experiments. From these isobaric labeling-based, global quantitative proteomic experiments, we were able to quantify a total of 12,295 proteins (FDR < 1%), with 5,169 proteins quantified across the entire panel of the SCLC cell lines (Supplementary Fig. 1F; Supplementary Table S2). Correlation analyses of the resulting protein abundances indicated that an excellent reproducibility was achieved in these quantitative proteomic experiments (Supplementary Fig. 1G).

We then performed a series of bioinformatic analyses to identify proteomic changes that might be associated with the selective sensitivity of certain SCLC cell lines to PARPi (Supplementary Fig. 1H). Specifically, we first grouped the SCLC cell lines and their respective proteomic datasets into four subtypes (i.e., ASCL1^high^, NEUROD1^high^, POU2F3^high^, and TF^low^). Because the vast majority of SCLC is characterized by the over-expression of ASCL1, we initially focused our analyses on the ASCL1^high^ SCLC cell lines. For these cell lines, we further divided them into the ASCL1^high^/talazoparib-sensitive (a total of 8 cell lines) and ASCL1^high^/talazoparib-resistant (a total of 6 cell lines) cell lines. We found that talazoparib was able to induce the downregulation of 256 proteins in the ASCL1^high^/talazoparib-sensitive cell lines. However, we did not identify any significantly downregulated proteins from the ASCL1^high^/talazoparib-resistant cell lines (Fig. 1C). We therefore termed the 256 down-regulated proteins as the PARPi-induced protein degradation signature (PiPS). Gene Ontology (GO) analyses showed that many of these PiPS proteins were involved in biological processes linked to the pathogenesis of SCLC, including regulation of transcription (*P* = 2.89 × 10^-6^), cell proliferation (*P* = 5.97 × 10^-6^), and cell cycle (*P* = 1.41 × 10^-5^) (Fig. 1D). Interestingly, several SCLC lineage-specific oncoproteins, including ASCL1, as well as two key jumonji histone demethylases (KDM4A and KDM5B) were among the PiPS, and these proteins were found to be down-regulated in response to talazoparib treatment (in ASCL1^high^/talazoparib-sensitive cell lines) (Supplementary Table S3).

Besides the ASCL1^high^ SCLC subtype, we also identified 10 and 85 PiPS proteins from the NEUROD1^high^ and POU2F3^high^ SCLC subtype, respectively (Supplementary Fig. 1I; Supplementary Table S3). Similar to ASCL1, we found that NEUROD1 and POU2F3 were also dramatically down-regulated in response to talazoparib treatment (in the respective talazoparib-sensitive cell lines) (Supplementary Table S3). Finally, because all the TF^low^ SCLC cell lines were resistant to talazoparib, we were not able to identify any PiPS proteins from this subtype (Supplementary Fig. 1I; Supplementary Table S3).

### SCLC Lineage-specific Oncoproteins are Degraded in Response to PARPi-induced Genotoxicity

The PiPS proteins contained several known SCLC lineage-specific oncoproteins (e.g., ASCL1, NEUROD1 and POU2F3). In addition, we also found, in PiPS (from the ASCL1^high^/talazoparib-sensitive subtype), several members of the histone demethylase family, including KDM4A (JMJD2A) and KDM5B (JARID1B). These two enzymes catalyze the removal of methyl groups from lysine residues (i.e., H3K9/H3K36, and H3K4, respectively), leading to the transcriptional regulation of gene expression^24^. Interestingly, these histone demethylases are known to be overexpressed in many types of cancers such as breast, prostate, and lung cancers and therefore have been proposed as potential oncoproteins^25–28^. Their functions in the pathogenesis of SCLC, however, are poorly understood.

We first validated our quantitative proteomic results and found that talazoparib treatment resulted in dramatic downregulation of ASCL1, KDM4A, and KDM5B in H2081 cells (Fig. 2A; Supplementary Fig. 2A-C). Talazoparib also reduced the levels of these proteins and tumor growth in an H2081 xenograft model (Fig. 2B and C; Supplementary Fig. 2D). We next performed quantitative real-time PCR (qRT-PCR) experiments and found that talazoparib treatment did not affect the mRNA levels of these proteins (Fig. 2D). These results also highlighted the power of quantitative proteomic in the identification of these post-translational regulation mechanisms.

**Fig. 2.**
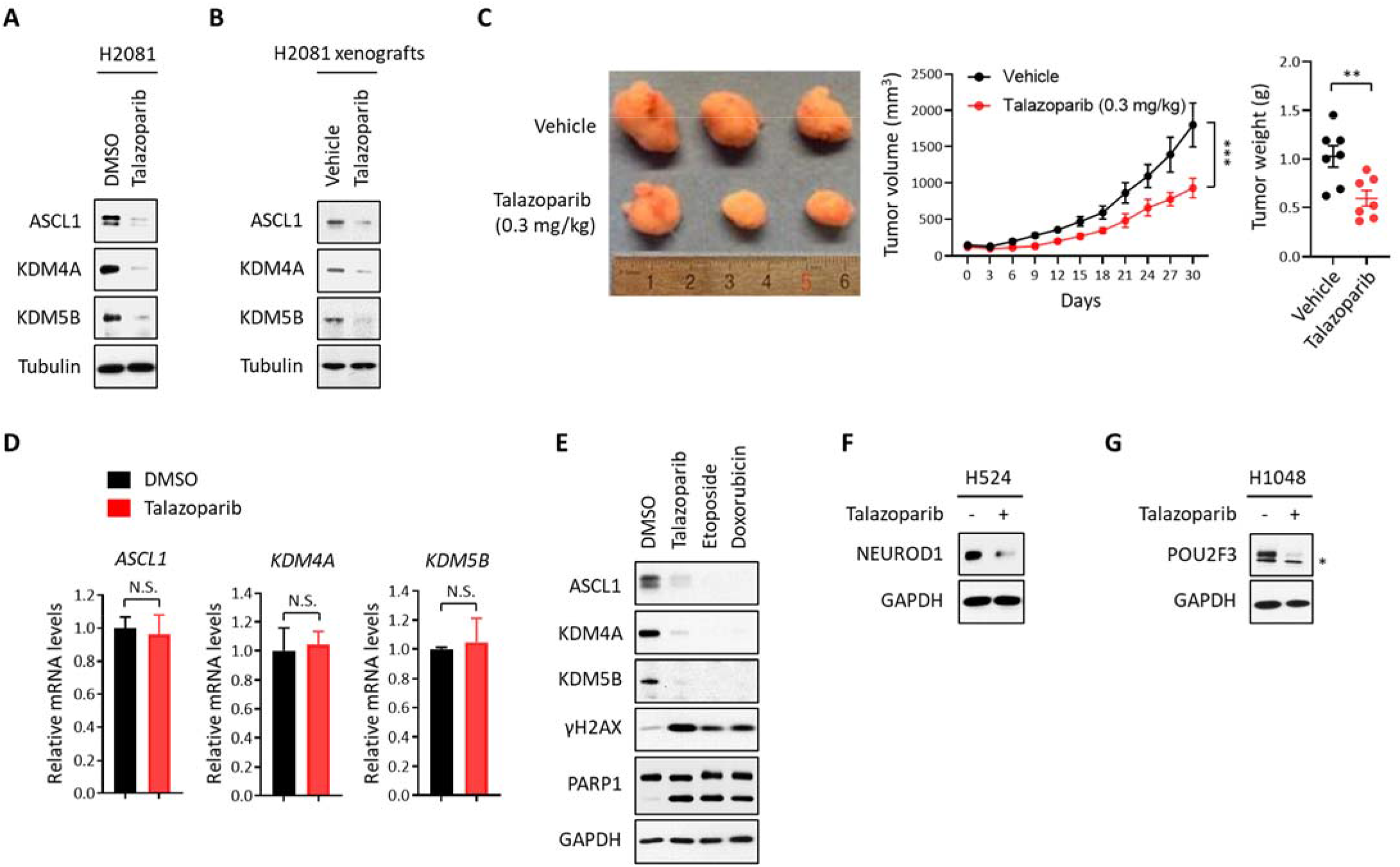
SCLC Lineage-specific Oncoproteins are Degraded in Response to PARPi-induced Genotoxicity. (A) The levels of ASCL1, KDM4A, and KDM5B in talazoparib treatment. H2081 cells were treated with or without talazoparib (1 μM for 48 hrs) and the cell lysates were subjected to immunoblot analysis using the indicated antibodies. See also Supplementary Fig. 2A for the quantification. (B) The levels of ASCL1, KDM4A, and KDM5B *in vivo*. H2081-implanted xenograft tumors were treated with or without talazoparib (0.3 mg/kg for 30 days) and the tumor extracts were subjected to immunoblot analysis using the indicated antibodies. See also Supplementary Fig. 2D for the quantification. (C) Toxicity of talazoparib *in vivo*. Mice implanted with H2081 cells were treated with or without talazoparib (0.3 mg/kg for 30 days). Left, the image of tumor size; Right, tumor volume and weight (n = 8–10 per each group). (D) The mRNA levels of ASCL1, KDM4A, and KDM5B in talazoparib treatment. H2081 cells were treated with or without talazoparib (1 μM for 48 hrs) and the mRNA levels of *ASCL1, KDM4A* and *KDM5B* were measured by qRT-PCR analysis. (E) The levels of ASCL1, KDM4A, and KDM5B in the treatment of DNA damaging agents. H2081 cells treated with talazoparib (1 μM), etoposide (1 μM), or doxorubicin (1 μM) for 48 hrs and the cell lysates were subjected to immunoblot analysis using the indicated antibodies. See also Supplementary Fig. 2E for the quantification. (F) The levels of NEUROD1 in talazoparib treatment. H524 cells were treated with talazoparib (1 μM for 48 hrs) and cell lysates were subjected to immunoblot analysis using the indicated antibodies. See also Supplementary Fig. 2F for the quantification. (G) The levels of POU2F3 in talazoparib treatment. H1048 cells were treated with talazoparib (1 μM for 48 hrs) and the cell lysates were subjected to immunoblot analysis using the indicated antibodies. The asterisk indicates a non-specific band. See also Supplementary Fig. 2G for the quantification.

Furthermore, the treatment of H2081 cells using genotoxic agents (e.g., etoposide and doxorubicin) also resulted in DDR (as shown by increased γH2AX levels), cytotoxicity (as shown by the increase in PARP1 cleavage), as well as the downregulation of ASCL1, KDM4A, and KDM5B (Fig. 2E; Supplementary Fig. 2E). Besides ASCL1, we observed that other lineage-specific oncoproteins (i.e., NEUROD1 and POU2F3) were also dramatically downregulated by talazoparib treatment in talazoparib-sensitive H524 cells (NEUROD1^high^) and H1048 cells (POU2F3^high^), respectively (Fig. 2F and G; Supplementary Fig. 2F and G).

In summary, our data showed that the SCLC lineage-specific oncoproteins were degraded in response to PARPi-induced genotoxicity. Because of the critical roles of these proteins in the pathogenesis of SCLC, our data also raise the hypothesis that the induced degradation of these lineage-specific oncoproteins could be a mechanism that underlies the selective cytotoxic effects of PARPi (as well as other DNA damaging agents) in SCLC.

### HUWE1 Senses PARPi-induced Genotoxicity to Degrade ASCL1

Because ASCL1 regulates a transcription program that is critical for the survival and proliferation of SCLC^29, 30^, we first investigated if PARPi treatment affects the expression of ASCL1 target genes. Quantitative RT-PCR analyses confirmed that the mRNA levels of several representative ASCL1 transcription targets, including *MYCL1, RET, SOX2*, and *BCL2* were significantly down-regulated in talazoparib-treated cells (Fig. 3A). To directly demonstrate the relevance of ASCL1 loss in mediating the cytotoxicity of PARPi, we depleted ASCL1 using two independent short hairpin RNAs (shRNAs). SCLC cells with ASCL1 knock-down (KD, shASCL1 #1 and #2) showed reduced survival and increased cell death (PARP1 cleavage), compared to control cells (Supplementary Fig. 3A and B). In the Cancer Dependency Map project (DepMap; www.depmap.org)^31^, the knock-down of ASCL1 also led to the decreased growth in 25 human SCLC cell lines (Supplementary Fig. 3C). On the contrary, overexpression of ASCL1 (ASCL1-Myc) attenuated cell death induced by talazoparib treatment (Fig. 3B; Supplementary Fig. 3D).

**Fig. 3.**
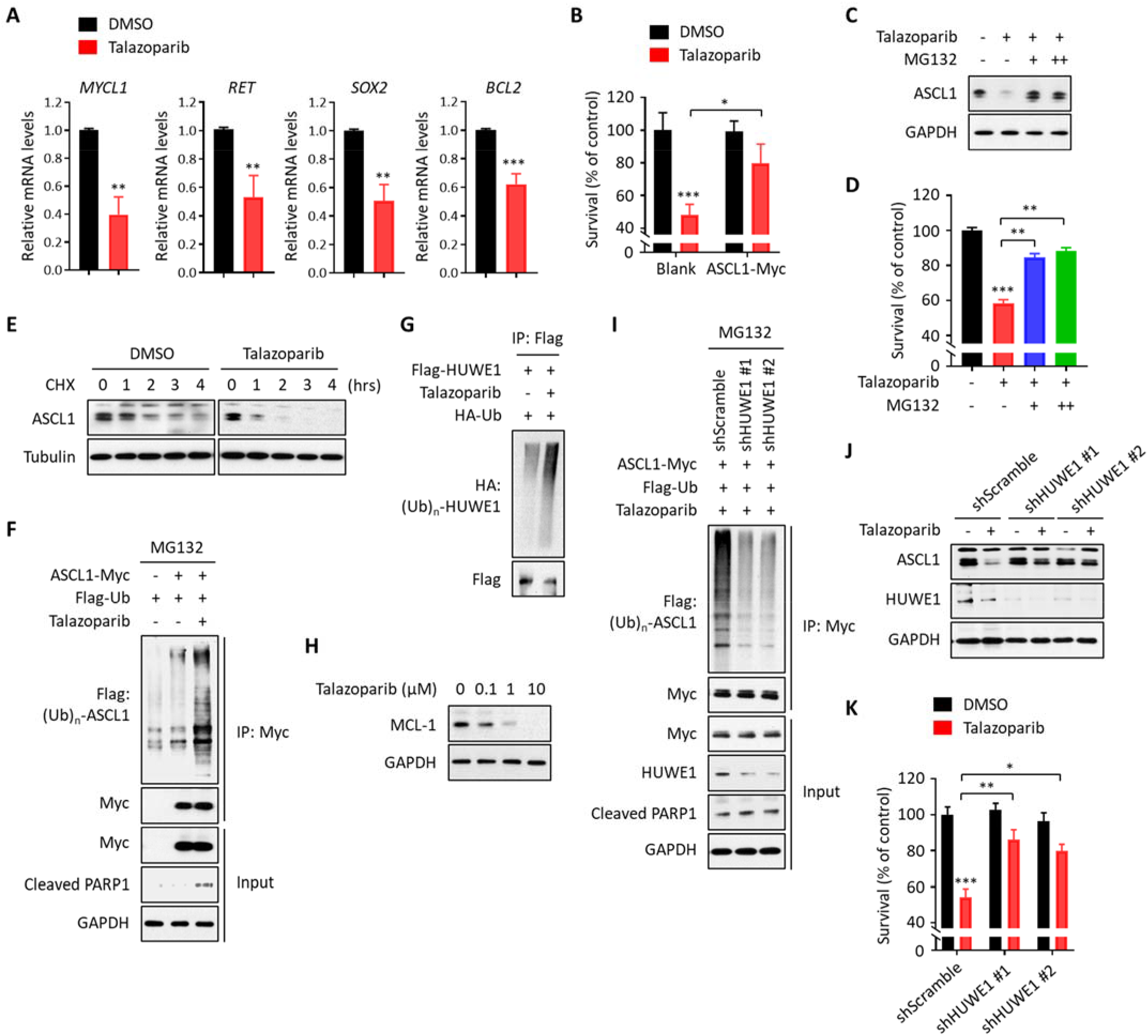
HUWE1 Senses PARPi-induced Genotoxicity to Degrade ASCL1. (A) qRT-PCR analysis of the downstream target genes of ASCL1. H2081 cells were treated with or without talazoparib (1 μM for 48 hrs) and the mRNA levels of the indicated target genes were measured by qRT-PCR analysis (n = 3). (B) Viability assays in ectopic expression of ASCL1. H2081 cells expressing ASCL1 (ASCL1-Myc) were treated with talazoparib (1 μM for 48 hrs) and viability was measured using a CellTiter-Glo assay (n = 3). (C) Inhibition of ASCL1 degradation. H2081 cells were treated with talazoparib (1 μM for 48 hrs) and then, proteasome inhibitor (MG132, 10 (+) and 20 (++) μM) was further treated for last 6 hrs. See also Supplementary Fig. 3E for the quantification. (D) Inhibition of cell death by proteasome inhibition. H2081 cells were treated with talazoparib (1 μM for 48 hrs) and then, MG132 (10 (+) and 20 (++) μM) was further treated for last 6 hrs (n = 3). Viability was measured using a CellTiter-Glo assay (n = 3). (E) Half-life of ASCL1. H2081 cells pre-treated with or without talazoparib (1 μM) were treated with cycloheximide (CHX, 10 μg/ml) as indicated. The cell lysates were subjected to immunoblot analysis using the indicated antibodies. See also Supplementary Fig. 3G for the quantification. (F) Ubiquitination of ASCL1. H2081 cells expressing ASCL1 (ASCL1-Myc) were treated with or without talazoparib (1 μM for 48 hrs) and then, MG132 (10 μM) was further treated for last 6 hrs. The cell lysates were subjected to immunoprecipitation analysis using anti-Myc antibody. Immunoprecipitates or inputs were resolved by SDS-PAGE and subjected to immunoblot analysis using the indicated antibodies. (G) Auto-ubiquitination of HUWE1 in talazoparib treatment. H2081 cells transfected with Flag-HUWE1 were treated with talazoparib (1 μM for 48 hrs) and the cell lysates were subjected to immunoprecipitation using anti-Flag M2 affinity gel beads. Immunoprecipitates were resolved by SDS-PAGE and subjected to immunoblot analysis using the indicated antibodies. (H) Downregulation of MCL-1 in talazoparib treatment. H2081 cells were treated with talazoparib for 48 hrs and the cell lysates were subjected to immunoblot analysis using the indicated antibodies. See also Supplementary Fig. 3M for the quantification. (I) Inhibition of ASCL1 ubiquitination. H2081 cells expressing ASCL1 (ASCL1-Myc) were depleted with HUWE1 (shHUWE1 #1 and #2) and treated with talazoparib (1 μM for 48 hrs). MG132 (10 μM) was further treated for last 6 hrs and the cell lysates were subjected to immunoprecipitation analysis using anti-Myc antibody. Immunoprecipitates or inputs were resolved by SDS-PAGE and subjected to immunoblot analysis using the indicated antibodies. (J) Inhibition of ASCL1 degradation. H2081 cells depleted with HUWE1 (shHUWE1 #1 and #2) were treated with talazoparib (1 μM for 48 hrs). The cell lysates were subjected to immunoblot analysis using the indicated antibodies. See also Supplementary Fig. 3N for the quantification. (K) Inhibition of cell death in depletion of HUWE1. H2081 cells depleted with HUWE1 (shHUWE1 #1 and #2) were treated with talazoparib (1 μM for 48 hrs). Viability was measured using a CellTiter-Glo assay (n = 3).

We next sought to determine the molecular mechanism of PARPi-induced ASCL1 downregulation. Intriguingly, we found that talazoparib-induced ASLC1 downregulation was completely blocked by a proteasome inhibitor, MG132, but not a pan-caspase inhibitor, Z-VAD (Fig. 3C; Supplementary Fig. 3E and F). Importantly, cell death was greatly reduced in treatment with both talazoparib and MG132 (Fig. 3D). We used cycloheximide (CHX) to block the synthesis of new proteins and found that talazoparib treatment led to accelerated turnover of ASCL1 (Fig. 3E; Supplementary Fig. 3G). Consistently, we also observed that ASCL1 ubiquitination levels were dramatically increased in response to talazoparib treatment, compared to control (Fig. 3F). Collectively, these data provide the evidence supporting a model where PARPi leads to the UPS-dependent degradation of ASCL1, and subsequently, cell death of SCLC.

Because PARPi causes PARP1 trapping, which, in return, leads to DDR^3, 4^, we hypothesized that an E3 ubiquitin ligase, whose activity is increased during DDR, could ubiquitinate ASCL1 and mediates its degradation. HUWE1 (HECT, UBA, and WWE domain-containing 1) is a HECT domain-containing E3 ubiquitin ligase that regulates many biological processes linked to DDR^32–36^. More importantly, its E3 ubiquitin ligase activity is known to increase upon sensing genotoxic stress^34–36^. In addition, a recent study showed that ASCL1 is targeted by HUWE1 for degradation in neural stem cells^37^. Whether HUWE1 mediates the degradation of ASCL1 during PARPi-induced DDR in SCLC is unknown. To study the potential association between ASCL1 and HUWE1, we performed co-immunoprecipitation assays and observed that ASCL1 interacted with HUWE1 (Supplementary Fig. 3H). Ectopic expression of HUWE1 led to the degradation of ASCL1 (Supplementary Fig. 3I), and this effect was blocked by MG132 (Supplementary Fig. 3J). Furthermore, the E3 ubiquitin ligase-dead (LD) mutant of HUWE1 did not degrade ASCL1 (Supplementary Fig. 3K). Finally, overexpression of HUWE1 enhanced ASCL1 ubiquitination (Supplementary Fig. 3L).

We next interrogated HUWE1-mediated ASCL1 degradation in the context of PARPi treatment. Immunoblotting analyses showed increased auto-ubiquitination of HUWE1 (Fig. 3G) and the degradation of known HUWE1 targets (i.e., MCL-1)^38^ in talazoparib-treated cells (Fig. 3H; Supplementary Fig. 3M). These results suggest that that PARPi-induced genotoxicity is able to activate the E3 ubiquitin ligase activity of HUWE1. Moreover, we found that talazoparib-induced ubiquitination and degradation of ASCL1 were dramatically reduced in HUWE1 KD (shHUWE1 #1 and #2) compared to control (Fig. 3I and J; Supplementary Fig. 3N). Furthermore, HUWE1 KD also greatly decreased the levels of talazoparib-induced cell death (Fig. 3K). Collectively, these data provide evidence that PARPi-induced genotoxicity activates HUWE1, which targets the degradation of ASCL1, and promotes the cell death of ASCL1^high^ SCLC.

### RNF8 Senses PARPi-induced Genotoxicity to Degrade KDM4A

Besides ASCL1, the PiPS proteins also contained several members of the histone demethylase family, including KDM4A and KDM5B (Fig. 2). Histone demethylase KDM4A is a member of the Jumonji domain 2 (JMJD2) family and its role is to catalyze the conversion of trimethylated histone (e.g., H3K9me3 and H3K36me3) to their corresponding demethylated forms^39, 40^. KDM4A is overexpressed in many cancer types of human malignancies including prostate, breast, lung, colorectal, and lymphoma and it has therefore been proposed as an oncogene^25, 39^. However, the role of KDM4A in the oncogenesis of SCLC and in its therapeutic response to PARPi is poorly understood.

We first validated our quantitative proteomics results and found that talazoparib treatment led to the downregulation of KDM4A, with a concomitant increase of H3K9me3 and H3K36me3 levels (Fig. 4A; Supplementary Fig. 4A). Because the H3K9me3 and H3K36me3 marks are primarily associated with transcriptional repression^41–43^, we also investigated if the PARPi-induced KDM4A downregulation would affect gene transcription in SCLC. Quantitative RT-PCR analyses showed that the mRNA expression levels of several representative KDM4A target genes including *HK2, PLK1*, and *TBL1X^44^* were down-regulated in talazoparib treatment (Fig. 4B). Next, we depleted KDM4A using two independent shRNAs (shKDM4A #1 and #2) and found that these cells had increased cell death (Supplementary Fig. 4B and C). In the Cancer DepMap project, KDM4A KD also led to decreased growth in 25 human SCLC cell lines (Supplementary Fig. 4D). Furthermore, in the xenograft mouse model, the KDM4A inhibitor, ML324, increased the level of both H3K9me3 and H3K36me3 (Fig. 4C; Supplementary Fig. 4E) and inhibited tumor growth (Fig. 4D). These results indicate that the downregulation of KDM4A is sufficient to drive the cell death of SCLC. On the contrary, the ectopic expression of KDM4A protected cells against talazoparib-induced cell death (Fig. 4E; Supplementary Fig. 4F). Taken together, these data indicate that the cytotoxic effects of PARPi in SCLC are also mediated, at least in part, by the downregulation of KDM4A.

**Fig. 4.**
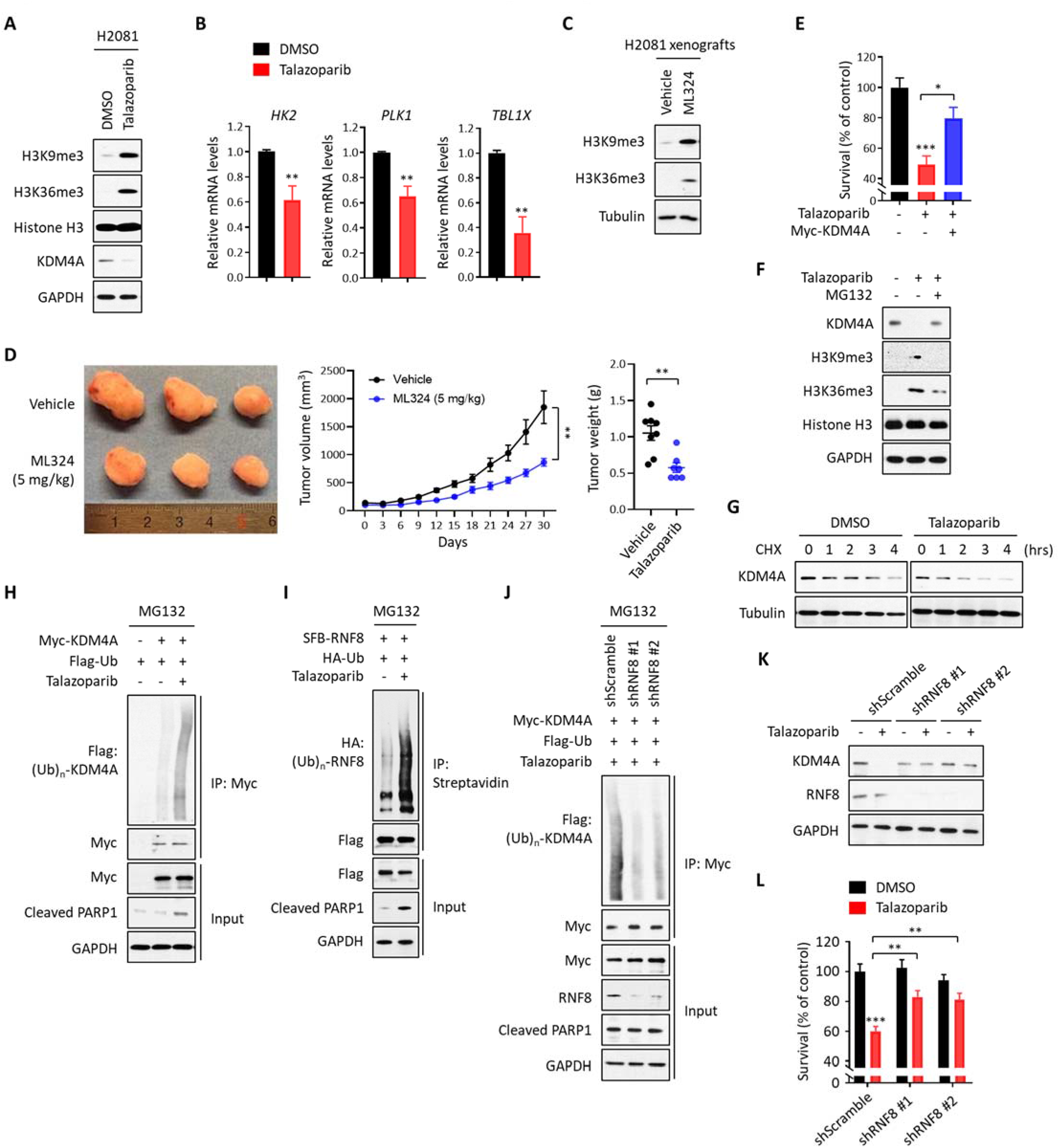
RNF8 Senses PARPi-induced Genotoxicity to Degrade KDM4A. (A) The levels of histone trimethylation (H3K9me3 and H3K36me3) in talazoparib treatment. H2081 cells were treated with or without talazoparib (1 μM for 48 hrs). The cell lysates were subjected to immunoblot analysis using the indicated antibodies. See also Supplementary Fig. 4A for the quantification. (B) qRT-PCR analysis of downstream target genes of KDM4A. H2081 cells were treated with or without talazoparib (1 μM for 48 hrs) and the mRNA levels of the indicated target genes were measured by qRT-PCR analysis (n = 3). (C) The levels of histone trimethylation (H3K9me3 and H3K36me3) *in vivo*. H2081-implanted xenograft tumors were treated with or without ML324 (5 mg/kg for 30 days) and the tumor extracts were subjected to immunoblot analysis using the indicated antibodies. See also Supplementary Fig. 4E for the quantification. (D) Toxicity of ML324 *in vivo*. Mice implanted with H2081 cells were treated with or without ML324 (5 mg/kg for 30 days). Left, the image of tumor size; Right, tumor volume and weight (n = 8–10 per each group). (E) Viability assays in ectopic expression of KDM4A. H2081 cells expressing KDM4A (Myc-KDM4A) were treated with talazoparib (1 μM for 48 hrs) and viability was measured using a CellTiter-Glo assay (n = 3). (F) The levels of KDM4A degradation and its subsequent histone trimethylation in proteasome inhibition. H2081 cells were treated with talazoparib (1 μM for 48 hrs) and then, MG132 (10 μM) was further treated for last 6 hrs. See also Supplementary Fig. 4G for the quantification. (G) The half-life of KDM4A. H2081 cells pre-treated with or without talazoparib (1 μM) were treated with CHX (10 μg/ml) as indicated. The cell lysates were subjected to immunoblot analysis using the indicated antibodies. See also Supplementary Fig. 4H for the quantification. (H) Ubiquitination of KDM4A. H2081 cells expressing KDM4A (Myc-KDM4A) were treated with or without talazoparib (1 μM for 48 hrs) and then, MG132 (10 μM) was further treated for last 6 hrs. The cell lysates were subjected to immunoprecipitation analysis using anti-Myc antibody. Immunoprecipitates or inputs were resolved by SDS-PAGE and subjected to immunoblot analysis using the indicated antibodies. (I) Auto-ubiquitination of RNF8 in talazoparib treatment. H2081 cells transfected with SFB-RNF8 were treated with or without talazoparib (1 μM for 48 hrs) and the cell lysates were subjected to immunoprecipitation using streptavidin beads. Immunoprecipitates or inputs were resolved by SDS-PAGE and subjected to immunoblot analysis using the indicated antibodies. (J) Inhibition of KDM4A ubiquitination. H2081 cells expressing KDM4A (Myc-KDM4A) were depleted with RNF8 (shRNF8 #1 and #2) and treated with talazoparib (1 μM for 48 hrs). MG132 (10 μM) was further treated for last 6 hrs and the cell lysates were subjected to immunoprecipitation analysis using anti-Myc antibody. Immunoprecipitates or inputs were resolved by SDS-PAGE and subjected to immunoblot analysis using the indicated antibodies. (K) Inhibition of KDM4A degradation. H2081 cells depleted with RNF8 (shRNF8 #1 and #2) were treated with talazoparib (1 μM for 48 hrs). The cell lysates were subjected to immunoblot analysis using the indicated antibodies. See also Supplementary Fig. 4M for the quantification. (L) Inhibition of cell death in depletion of RNF8. H2081 cells depleted with RNF8 (shRNF8 #1 and #2) were treated with talazoparib (1 μM for 48 hrs). Viability was measured using a CellTiter-Glo assay (n = 3).

We next investigated the molecular mechanism by which PARPi regulates the downregulation of KDM4A. Because talazoparib treatment led to the downregulation of KDM4A protein levels, but not its mRNA (Fig. 2A and D), we investigated the role of UPS in KDM4A regulation. Interestingly, talazoparib-induced KDM4A downregulation, as well as the increase in H3K9me3 and H3K36me3 levels, were largely blocked by MG132 treatment (Fig. 4F; Supplementary Fig. 4G). In addition, the turnover of KDM4A in CHX-treated cells was dramatically accelerated after talazoparib treatment (Fig. 4G; Supplementary Fig. 4H). Consistent with this model, KDM4A ubiquitination levels also were dramatically increased in talazoparib-treated cells (Fig. 4H). Collectively, these data suggest that PARPi may lead to the degradation of KDM4A via UPS.

Based on these data, we hypothesized that, similar to ASCL1, KDM4A could also be degraded by an E3 ubiquitin ligase that is activated during DDR. RNF8 is an E3 ubiquitin ligase that plays a critical role in maintaining genomic stability by promoting the repair of DSB through ubiquitin signaling^45–49^. Although RNF8 is known to recruit RNF168 as well as downstream repair factors (e.g., 53BP1 and *BRCA1/2*) to sites of DSB on K63-linked ubiquitination, accumulating data have also shown that RNF8 could degrade its target proteins by promoting K48-linked ubiquitination^50, 51^. Indeed, a recent study demonstrated that the RNF8-dependent degradation of KDM4A triggers 53BP1 recruitment to DNA damage sites^52^. These raise an interesting hypothesis that RNF8 could function as an E3 ubiquitin ligase that mediates the PARPi-induced degradation of KDM4A in SCLC. To directly test this hypothesis, we first carried out co-immunoprecipitation assays and found that KDM4A interacted with RNF8 (Supplementary Fig. 4I). Ectopic expression of RNF8 led to the degradation of KDM4A (Supplementary Fig. 4J), an effect that could be completely blocked by MG132 (Supplementary Fig. 4K). Furthermore, the ectopic expression of RNF8 dramatically increased KDM4A ubiquitination (Supplementary Fig. 4L). Taken together, these data showed that KDM4A is a substrate of RNF8, and it is ubiquitinated and degraded by RNF8 in SCLC.

We next examined the role of RNF8 in PARPi-induced KDM4A degradation. First, we observed that talazoparib treatment strongly activated the E3 ubiquitin ligase activity of RNF8, as shown by the profound increase in the auto-ubiquitination levels of RNF8 (Fig. 4I). Talazoparib-induced ubiquitination of KDM4A was dramatically decreased in RNF8-depleted cells, compared to control (Fig. 4J). Furthermore, talazoparib-induced KDM4A degradation was completely blocked in RNF8-depleted cells (Fig. 4K; Supplementary Fig. 4M). RNF8 depletion was also able to protect cells against talazoparib-induced cell death (Fig. 4L). Collectively, these results indicate that RNF8 is the major E3 ligase for KDM4A ubiquitination and degradation during PARPi-induced genotoxicity in SCLC.

In addition to KDM4A, we also identified another member of the histone demethylases KDM5B^28, 53^ as one of the down-regulated proteins after talazoparib treatment (Fig. 2). Accordingly, we found that talazoparib treatment increased the level of H3K4me3 in SCLC (Supplementary Fig. 4N). Similar to KDM4A, KDM5B promotes the cell survival of SCLC in talazoparib treatment (Supplementary Fig. 4O-S) and talazoparib-induced KDM5B degradation was also mediated by UPS (Supplementary Fig. 4T-V). Unlike KDM4A, however, the E3 ubiquitin ligase responsible for KDM5B degradation is currently unknown, which warrants further investigation.

Collectively, these data indicate that the PARPi treatment in SCLC induces genotoxic stress, which leads to the activation of certain E3 ubiquitin ligases (e.g., HUWE1 and RNF8), and subsequently, ubiquitination and degradation of key survival proteins for SCLC (e.g., ASCL1 and KDM4A).

### The Synergistic Effects Between PARPi and PiPS-targeting Agents

While chemotherapy often results in significant response and tumor shrinkage in SCLC, relapse is almost universal^12, 13^. Because the downregulation of PiPS proteins serves as a major therapeutic response to PARPi in SCLC, we tested whether co-targeting these PiPS proteins and PARP1 could offer a therapeutic strategy to achieve a more complete and durable response. Toward this, we used JQ-1 to reduce ASCL1 expression^54^, ML324 to inhibit KDM4A^55^, and PBIT to inhibit KDM5B^56, 57^ (Fig. 5A; Supplementary Fig. 5H). Consistent with previous findings, ASCL1 expression was dramatically reduced in ASCL1^high^ H2081 cells and xenograft tumors treated with JQ-1 (Fig. 5B; Supplementary Fig. 5A). ML324 and PBIT treatment also led to a considerable increase of H3K9me3, H3K36me3, and H3K4me3, respectively (Fig. 5B and 4C; Supplementary Fig. 5I). Indeed, these compounds all significantly reduced the survival in H2081 cells (Supplementary Fig. 5B) and xenograft tumor growth (Supplementary Fig. 5C; Fig. 4D).

**Fig. 5.**
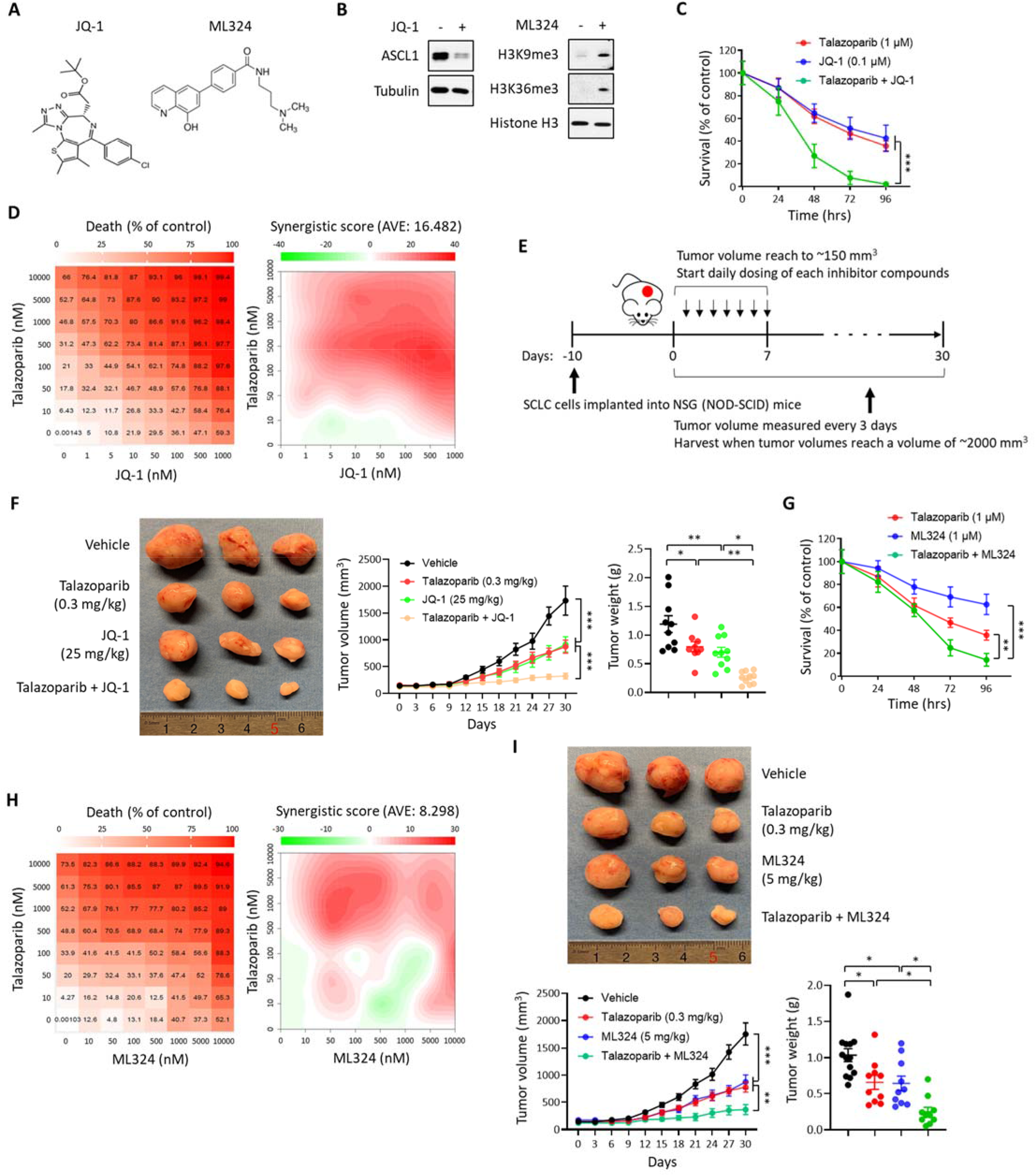
The Synergistic Effects Between PARPi and PiPS-targeting Agents. (A) The structure of the small-molecule compounds JQ-1 and ML324. (B) The effect of small-molecule compounds JQ-1 and ML324. H2081 cells were treated with JQ-1 (0.1 μM for 48 hrs) or ML324 (1 μM for 48 hrs) and the cell lysates were subjected to immunoblot analysis using the indicated antibodies. (C) The synergistic effect of talazoparib with JQ-1 in a time-dependent manner. H2081 cells treated with talazoparib (1 μM), JQ-1 (0.1 μM), or talazoparib + JQ-1 in a time-dependent manner. Viability was measured using a CellTiter-Glo assay (n = 3). (D) The synergistic effect of talazoparib with JQ-1 in a concentration-dependent manner. H2081 cells treated with talazoparib, JQ-1, or talazoparib + JQ-1 in a concentration-dependent manner. Viability was measured using a CellTiter-Glo assay (n = 3). Left, cell death ratio; Right, synergistic score. (E) Overview of H2081-implanted xenograft tumor models. (F) The synergistic effects of talazoparib with JQ-1 *in vivo*. Mice implanted with H2081 cells were treated with talazoparib (0.3 mg/kg), JQ-1 (25 mg/kg), or talazoparib + JQ-1 as indicated (n = 8-10 per each group). Left, the image of tumor size; Right, tumor volume and weight. (G) The synergistic effect of talazoparib with ML324 in a time-dependent manner. H2081 cells treated with talazoparib (1 μM), ML324 (1 μM), or talazoparib + ML324 in a time-dependent manner. Viability was measured using a CellTiter-Glo assay (n = 3). (H) The synergistic effect of talazoparib with ML324 in a concentration-dependent manner. H2081 cells treated with talazoparib, ML324, or talazoparib + ML324 in a concentration-dependent manner. Viability was measured using a CellTiter-Glo assay (n = 3). Left, cell death ratio; Right, synergistic score. (I) The synergistic effects of talazoparib with ML324 *in vivo*. Mice implanted with H2081 cells were treated with talazoparib (0.3 mg/kg), ML324 (5 mg/kg), or talazoparib + ML324 as indicated (n = 8-10 per each group). Top, the image of tumor size; Bottom, tumor volume and weight.

We next evaluated the synergistic effect of PARPi in combination with the aforementioned inhibitors in SCLC. Indeed, the survival and tumor growth were more dramatically reduced in H2081 cells and xenograft tumors treated with a combination of talazoparib and JQ-1, compared to talazoparib or JQ-1 alone (Fig. 5C-F; Supplementary Fig. 5D). Similarly, a combination of talazoparib with ML324 also led to profound cell death and the inhibition of tumor growth compared to talazoparib or ML324 alone (Fig. 5G-I; Supplementary Fig. 5E). Mice that received single or the combination treatment in xenograft experiments did not show any significant changes in body weight (Supplementary Fig. 5F and G). Similar synergistic effects were also observed between talazoparib and PBIT (Supplementary Fig. 5J-L). We previously showed that a secreted protein, IGFBP5, is a transcription target of ASCL1^58^. Because IGFBP5 is an inhibitor of IGF signaling, downregulation of ASCL1 leads to the lower expression of IGFBP5, and hence, the paradoxical activation of IGF signaling. The regulation of IGF1R activity provides an alternative mechanism to affect cell survival under ASCL1-repressed conditions. Indeed, we found that the combination of talazoparib with an IGF1R inhibitor (BMS-754807) was highly synergistic (Supplementary Fig. 5M-R).

Taken together, these data show that co-targeting PARP1 and the other PiPS proteins including ASCL1, KDM4A, and KDM5B were able to elicit dramatic cytotoxic effects, compared to PARPi or PiPS-targeting agents alone. The full therapeutic potential of these combination strategies warrants further studies.

### Predictive Biomarkers for PARPi-mediated Cytotoxicity in SCLC

PARP1 has been proposed as a promising therapeutic target for SCLC^59, 60^. However, *BRCA1/2* mutations are rarely found in SCLC, and the mechanism for the selective PARPi toxicity in certain SCLC cell lines is poorly understood. Towards this, we selected representative SCLC cell lines from the talazoparib-sensitive group (i.e., H209, H1876, H2107, H2081, H128, H69, H1092, H2171, H524, and H1048), and also from the talazoparib-resistant group (i.e., H1436, H889, H82, H378, SW1271, and H1341) (Supplementary Fig. 6A-C). Despite the heterogeneous response of these SCLC cell lines to PARPi, we found that talazoparib treatment was able to elicit a general DDR in these cells, as shown by the universal increase in γH2AX levels (Fig. 6A; Supplementary Fig. 6D). These results suggest that the selective toxicity of PARPi in certain SCLC could result from a cell context-specific response to PARPi-induced DDR stimuli, rather than an unequal DDR signal itself.

**Fig. 6.**
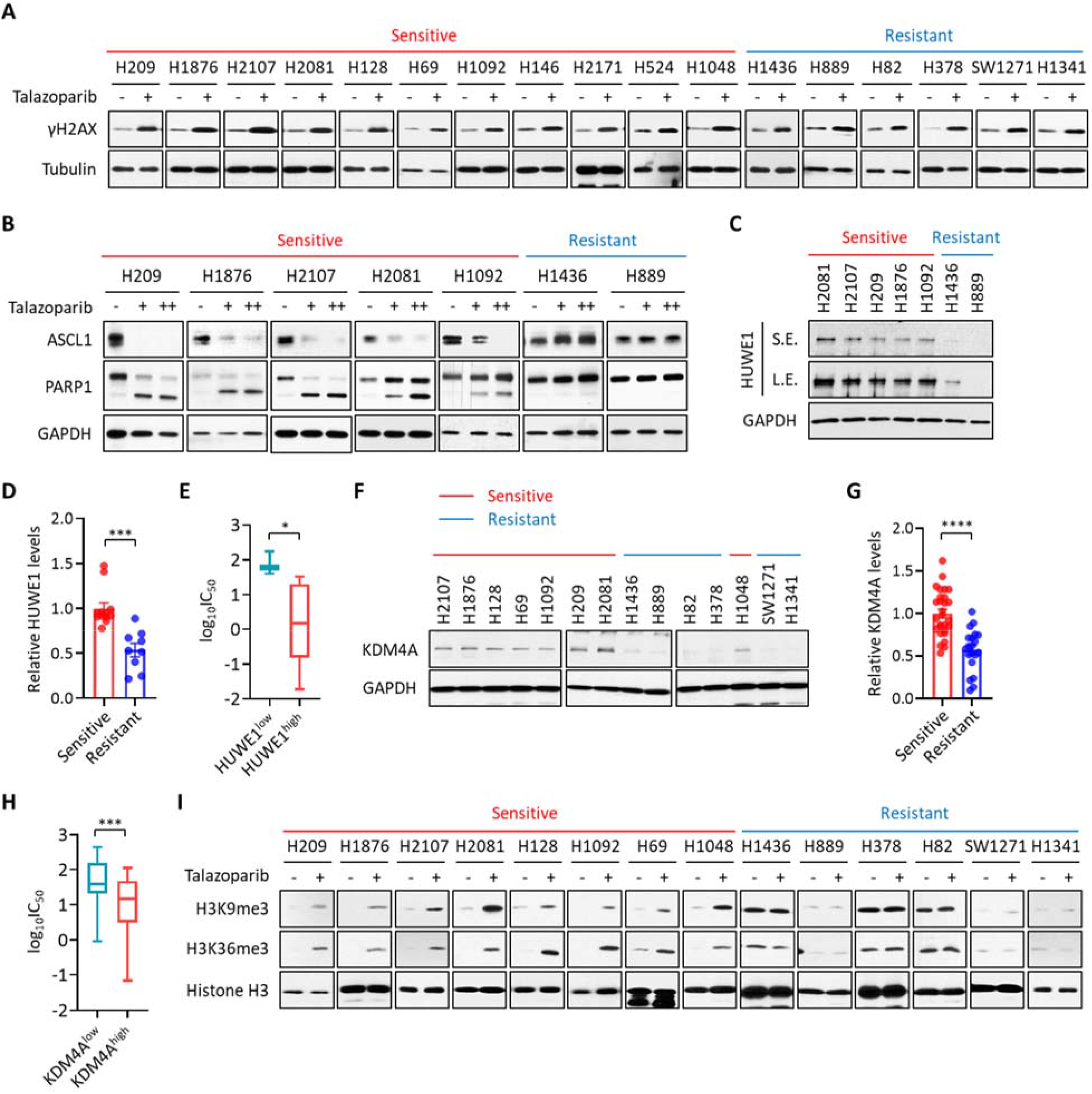
Predictive Biomarkers for PARPi-mediated Cytotoxicity in SCLC. (A) The extent of DNA damage in a panel of SCLC cell lines treated with talazoparib. A panel of SCLC cell lines were treated with talazoparib (1 μM for 48 hrs). The cell lysates were subjected to immunoblot analysis using the indicated antibodies. Red: sensitive SCLC cell lines; Blue: resistant SCLC cell lines. See also Supplementary Fig. 6D for the quantification. (B) The levels of ASCL1 and cleaved PARP1 in a panel of SCLC cell lines treated with talazoparib. A panel of SCLC cell lines were treated with talazoparib (1 (+), 10 (++) μM for 48 hrs). The cell lysates were subjected to immunoblot analysis using the indicated antibodies. Red: sensitive SCLC cell lines; Blue: resistant SCLC cell lines. See also Supplementary Fig. 6E for the quantification. (C) The levels of HUWE1 in a panel of ASCL1^high^ SCLC cell lines. Red: sensitive SCLC cell lines; Blue: resistant SCLC cell lines. See also Supplementary Fig. 6J for the quantification. (D) The relative levels of HUWE1 in ASCL1^high^ SCLC cell lines from TMT-based proteomic datasets. The values of HUWE1 from the DMSO-treated samples in PARPi-sensitive and -resistant ASCL1^high^ SCLC cell lines were collected and normalized. See also Supplementary Table S2. (E) Boxplots showing the sensitivity of a panel of ASCL1^high^ SCLC cell lines to PARPi. ASCL1^high^ SCLC cell lines were grouped by the relative levels of HUWE1 expression. The sensitivity was determined by the analysis of the GDSC database measuring IC_50_ against talazoparib treatment. (F) The levels of KDM4A in a panel of SCLC cell lines. Red: sensitive SCLC cells; Blue: resistant SCLC cell lines. See also Supplementary Fig. 6K for the quantification. (G) The relative levels of KDM4A in a panel of SCLC cell lines from TMT-based proteomic datasets. The values of KDM4A from the DMSO-treated samples in PARPi-sensitive and -resistant SCLC cell lines were collected and normalized. See also Supplementary Table S2. (H) Boxplots showing the sensitivity of a panel of SCLC cell lines to PARPi. SCLC cell lines were grouped by the relative levels of KDM4A expression. The sensitivity was determined by the analysis of the GDSC database measuring IC_50_ against talazoparib treatment. (I) Histone H3 trimethylation in a panel of SCLC cells treated with talazoparib. A panel of SCLC cell lines were treated with talazoparib (1 μM for 48 hrs) and the cell lysates were subjected to immunoblot analysis using the indicated antibodies. Red: sensitive SCLC cell lines; Blue: resistant SCLC cell lines. See also Supplementary Fig. 6M for the quantification.

From the aforementioned cell lines (both the talazoparib-sensitive and the talazoparib-resistant ones), we further examined the ones that expressed ASCL1 (i.e., ASCL1^high^, H209, H1876, H2107, H2081, H1092, H1436, and H889). We treated these cells with talazoparib and observed that the degree of ASCL1 degradation was highly correlated with the level of talazoparib toxicity (Fig. 6B). Specifically, talazoparib treatment led to increased ASCL1 degradation in ASCL1^high^/talazoparib-sensitive cells, but not in ASCL1^high^/talazoparib-resistant cells. In addition, similar results were obtained when these cells were treated with talazoparib in a dose-dependent manner (Supplementary Fig. 6F-I). Besides ASCL1, we also observed the degradation of NEUROD1 in NEUROD1^high^/talazoparib-sensitive cells, but not in NEUROD1^high^/talazoparib-resistant cells (Supplementary Fig. 6E). We observed robust degradation of POU2F3 in H1048 cells, which is a POU2F3^high^/talazoparib-sensitive cell line. Finally, we examined two TF^low^ cell lines (SW1271 and H1341) and found both to be resistant to talazoparib. These results point to that the degradation of lineage-specific transcription factors correlated with PARPi sensitivity in SCLC cell lines.

Since HUWE1 plays a critical role in regulating the PARPi-induced degradation of ASCL1 (Fig. 3), we hypothesized that HUWE1 could be a predictive biomarker for PARPi. Towards this, we first measured the abundance of HUWE1 in a representative panel of ASCL1^high^ SCLC cell lines. We observed that the PARPi-sensitive ASCL1^high^ SCLC cell lines expressed higher levels of HUWE1, compared to PARPi-resistant ASCL1^high^ SCLC cell lines (Fig. 6C; Supplementary Fig. 6J). Using TMT-based quantitative mass spectrometric analyses (Fig. 1), we performed proteomic profiling experiments of 24 SCLC cell lines treated with either DMSO or talazoparib. From this dataset, we extracted the basal HUWE1 abundances from the ASCL1^high^ SCLC cell lines (under the DMSO-treated conditions). From these analyses, we also showed that the ASCL1^high^/talazoparib-sensitive cells had higher HUWE1 levels, compared to the ASCL1^high^/talazoparib-resistant cells (Fig. 6D). To further investigate whether HUWE1 expression had an impact on the sensitivity of ASCL1^high^ SCLC cells to PARPi, we analyzed the drug response of ASCL1^high^ SCLC cell lines from the Genomics of Drug Sensitivity in Cancer database (GDSC, https://www.cancerrxgene.org/)^61^. The expression of HUWE1 in these ASCL1^high^ SCLC cell lines was obtained from a publicly available database^62^. We observed that ASCL1^high^ SCLC cell lines expressing higher levels of HUWE1 were more sensitive to talazoparib (Fig. 6E). Taken together, these data indicate that HUWE1 mediates the PARPi-dependent degradation of ASCL1, and that HUWE1 abundances could be a factor that determines the sensitivity of ASCL1^high^ SCLC cells to PARPi.

In addition to HUWE1, we also investigated whether KDM4A could function as another predictive biomarker for PARPi. We first measured the level of KDM4A in a representative panel of SCLC cell lines. We observed that PARPi-sensitive SCLC cell lines (i.e., H2107, H1876, H128, H69, H1092, H209, H2081, and H1048) expressed higher levels of KDM4A, compared to PARPi-resistant SCLC cell lines (i.e., H1436, H889, H82, H378, SW1271 and H1341) (Fig. 6F; Supplementary Fig. 6K). These results were further validated using our TMT-based proteomic dataset (from the DMSO-treated SCLC cell lines) (Fig. 6G). Analyses of PARPi sensitivity of SCLC cell lines in the GDSC database and the gene expression of SCLC lines also showed that SCLC cell lines with higher expression of KDM4A were significantly more sensitive to talazoparib (Fig. 6H). These results suggest that the abundance of KDM4A could also determine the sensitivity of SCLC cells to PARPi. We treated the aforementioned SCLC cell lines with talazoparib and observed robust KDM4A degradation in PARPi-sensitive SCLC cell lines (i.e., KDM4A^high^ SCLC cell lines, including H209, H1876, H2107, H2081, H128, H69, H1092, and H1048) (Supplementary Fig. 6L). Consistent with the degradation of KDM4A, H3K9me3 and H3K36me3 levels were dramatically increased in PARPi-sensitive SCLC cell lines, but not in PARPi-resistant cells, in response to talazoparib treatment (Fig. 6I; Supplementary Fig. 6M). These data point to the possibility that the abundance of KDM4A (and its degradation) could also be a factor that determines the selective cytotoxicity of PARPi in SCLC.

## Discussion

The critical roles of PARP1 in mediating DDR provide the rationale for the usage of PARPi to treat human malignancy. In particular, it is well known that cancers with homologous recombination deficiencies (HRD, as a result, for example *BRCA1/2* mutations) are particularly sensitive to PARPi, as a result of the synthetic lethality mechanism^7^. In addition to *BRCA1/2-mutated* breast and ovarian cancers, recent studies have indicated that a number of other human malignancies could also benefit from PARPi. For example, it is known that at least a subset of SCLC responds favorably to PAPRi, and based on these results, multiple clinical trials have been initiated for the evaluation of PARPi in SCLC^63^. However, the underlying mechanism(s) of PARPi sensitivity in SCLC is poorly understood. Mutations of *BRCA1/2* are not commonly found in this tumor type (e.g., less than 3% of the SCLC cases contain *BRCA1/2* mutations)^14, 15^. Using a series of PDX (patient-derived xenograft) models, a recent study evaluated whether HRD scores could predict PARPi sensitivity in SCLC^64^. In this study, no association was found between the *in vivo* sensitivity to talazoparib and the HRD score, or the mutation status of common DDR genes. These results point to a disease-specific context for the HRD to predict PARPi sensitivity. It was also concluded from this study that standard predictive biomarkers for PARPi in other cancer types (HRD scores, mutational burden and DDR gene mutations) are unlikely to be applicable in SCLC.

Although gene expression profiling experiments have been widely used to characterize tumor adaptive response to chemical perturbations, mRNA levels alone do not fully recapitulate these adaptive changes. Towards this, we used isobaric labeling-based, quantitative MS experiments to evaluate how the SCLC proteome is remodeled upon the treatment of PARPi. To do so, the global quantitative proteomic experiments were performed in a panel of 24 SCLC cell lines that were treated with talazoparib. These cells signify the common subtypes of SCLC, i.e., ASCL1^high^, NEUROD1^high^, POU2F3^high^, and low expression of all three transcription factors (TF^low^).

Because of the profound heterogeneity in the SCLC proteome and the limited number of the talazoparib-sensitive and talazoparib-resistant cell lines for each SCLC subtype (i.e., ASCL1^high^, NEUROD1^high^, POU2F3^high^ and TF^low^), the aggregation of the ratios (talazoparib vs. DMSO) across the cell lines and biological replicates appears to have caused the low average log2FC. Indeed, the comparison of the talazoparib-sensitive or talazoparib-resistant cell lines based on the log2FC was also less ideal to capture the proteomic changes associated with each subgroup. We hereby used a workflow where we first divided each subtype (e.g., ASCL1^high^ and NEUROD1^high^) into two categories based on their sensitivities to talazoparib treatment (i.e., talazoparib-sensitive and talazoparib-resistant cell lines) (Supplementary Fig. 1H). The protein abundances were directly extracted from the DMSO and talazoparib treatment samples for each cell line. In this step, we also calculated the *P* value and adjusted *P* value for every protein and identified a list of proteins that were significantly downregulated in either the talazoparib-sensitive or the talazoparib-resistant subgroup. Next, we performed cross-reference analyses to extract the proteins that were downregulated (in response to talazoparib treatment) only in the talazoparib-sensitive groups. Using this workflow, we discovered that ASCL1, KDM4A and KDM5B were downregulated only in the talazoparib-sensitive ASCL1^high^ groups. Using similar analyses, we also identified NEUROD1 and POU2F3 as talazoparib-induced downregulated proteins from the NEUROD1^high^ and POU2F3^high^ subtypes.

We believe that such analyses led to the identification of key proteomic changes that might explain talazoparib sensitivity in SCLC. As an example, ASCL1 is a transcription factor that is over-expressed in ~70% of the SCLC cases. ASCL1 is required for the establishment of pulmonary NE lineage and therefore is a lineage-specific oncoprotein for SCLC^21, 29^. Furthermore, it is required for the continued survival of SCLC *in vitro*, and for tumor formation in a genetically engineered mouse model of SCLC^65, 66^. Besides ASCL1, KDM4A and KDM5B are frequently over-expressed in various solid tumors and hence have been proposed as oncogenes.

We confirmed that talazoparib treatment down-regulates both transcription factors (e.g., ASCL1), and histone demethylases (e.g., KDM4A and KDM5B), indicating that these oncoproteins could be a critical mediator of the PARPi-induced cytotoxicity in SCLC. How does talazoparib regulate the abundance of these PiPS proteins? We found that talazoparib treatment did not alter the mRNA abundances of these proteins, suggesting that the PARPi-mediated regulation occurred at the post-transcriptional level. These results also highlighted the power of quantitative proteomics in identifying translational or post-translational regulation events. Ubiquitination is known to play a crucial role in regulating various forms of DDR. We performed biochemical characterization and showed that ASCL1 and KDM4A were degraded by two E3 ligases, HUWE1 and RNF8, respectively. Both HUWE1 and RNF8 are known to be induced in DDR to regulate genomic stability by promoting the repair of DSB through ubiquitin signaling^35, 36, 45–48^. HUWE1 has been previously proposed as a tumor suppressor^33, 67–69^. Specifically, HUWE1 KO results in increased oncogenesis in a mouse model of skin cancer. Consistent with its role as a tumor suppressor, besides ASCL1, several additional oncoproteins are also known substrates of HUWE1, including MYC and MCL-1. RNF8 is also an E3 ubiquitin ligase that plays a key role in the DNA damage response. Cells derived from RNF8 KO mice have increased radiosensitivity, class switch recombination (CSR) defects, and elevated genomic instability^70, 71^. Interestingly, RNF8 KO mice also demonstrated an increased risk for tumor development, suggesting that RNF8 could be a *bona fide* tumor suppressor^70^. Here we discovered that HUWE1 and RNF8 function as “sensors” to detect PARPi-induced DDR. The activation of these E3 ligases subsequently causes the degradation of ASCL1 and KDM4A, leading to the death of the corresponding SCLC cells. However, the E3 ubiquitin ligases that are involved in degrading the other lineage-specific oncoproteins (e.g., NEUROD1 and POU2F3), and the histone demethylase KDM5B are currently unknown.

Intriguingly, we also found that PARPi treatment caused the downregulation of several other UPS-related proteins, including ZFAND5 and UBE2T (Supplementary Fig. 7). For example, ZFAND5 is a member of the ubiquitous ZFAND protein family that binds to the 26S proteasome, and in doing so, stimulates its capacity to degrade the ubiquitinated proteins. It is conceivable that the downregulation of ZFAND5 could reduce the protein degradation via the 26S proteasome pathway. As another example, UBE2T is a ubiquitin-conjugating enzyme (E2) that is known to bind to FANCL, the ubiquitin E3 ligase component of the multiprotein Fanconi anemia (FA) complex. This molecular event is critical for the mono-ubiquitination of FANCD2, a key step in the FA DNA damage pathway. Therefore, the PARPi-induced downregulation of UBE2T also could reduce the ubiquitination of its target proteins (e.g., FANCD2), which serves as an independent mechanism to trigger additional DNA damage^72–74^. Indeed, recent CRISPR screens showed that the deletion of UBE2T sensitized the cells to olaparib treatment^75^. Whether these mechanisms (e.g., mediated by ZFAND5 and UBE2T) also contribute to the PARPi-induced cytotoxicity in SCLC warrants future studies.

Although SCLC usually responds to initial chemo- and/or radiotherapy approaches, resistance often rapidly develops^10, 12, 76, 77^. Thus, to develop the new combination therapies may inhibit the development of drug resistance, leading to more complete and durable control of the disease. Towards this, we also explored the combination of talazoparib with agents targeting the PiPS proteins known as oncoproteins to maximize the vulnerability of SCLC. Co-treatment of talazoparib and JQ-1 (targeting ASCL1) dramatically increased cell death both *in vitro* and *in vivo* (Fig. 5C-F; Supplementary Fig. 5D). In this case, ASCL1 protein levels were greatly reduced by two orthogonal mechanisms, i.e., JQ-1-mediated transcription reduction and PARPi-mediated ASCL1 degradation. Our previous study showed that JQ-1 treatment leads to the paradoxical activation of IGF signaling in SCLC^58^. Here, we also showed that PARPi also synergizes with IGF1R inhibitors (Supplementary Fig. 5P-R). Finally, a profound synergistic effect was also observed between PARPi and a KDM4A inhibitor ML324 or KDM5B inhibitor PBIT (Fig. 5G-I; Supplementary Fig. 5E and J-L). In this case, these two KDMs were also inhibited by two independent mechanisms, i.e., talazoparib-induced degradation and ML324- or PBIT-induced inhibition. The combination of PARPi with agents targeting these lineage-specific oncoproteins could offer a strategy to achieve a more complete and durable control of the disease. Finally, it has been suggested that the novel inflamed SCLC subtype (SCLC-I) expresses lower ASCL1, NEUROD1, and POU2F3, which might benefit from the addition of anti-PD-L1 to chemotherapy^78, 79^. Although our current study is focused on the SCLC subtypes that highly express ASCL1, NEUROD1, or POU2F3, we and others have recently shown that by triggering PARP1 trapping, PARP1 inhibitors induce the formation of cytosolic micronuclei, and thereby cause the activation of innate immune signaling^5, 80–82^. Our results therefore point to the exciting possibility of combining PARP1 inhibitors with immune checkpoint inhibitors in SCLC. The full therapeutic potential of these regimens will be investigated in future studies.

Talazoparib is a highly potent PARPi and its daily dosing is relatively low in patients. The distribution of PARPi *in vivo*, however, could vary dramatically, depending on the different anatomical sites. For example, using a radioactive version of olaparib (^18^F-olaparib) with PET (Positron Emission Tomography) imaging, Wilson et al. previously determined the tissue-specific distribution/accumulation of PARPi (i.e., olaparib) in animals^76^. The concentration of ^18^F-olaparib in the large intestine could be over 140-fold higher than that in the blood. Similarly, they also found that ^18^F-olaparib was rapidly taken up by and hence accumulated in xenograft tumors, suggesting that the intra-tumor concentrations of ^18^F-olaparib could be much higher, compared to those in the blood. The authors and others also demonstrated that the uptake of ^18^F-olaparib by tumors was based on its on-target engagement (i.e., binding to PARP1), and the level of ^18^F-olaparib accumulation was closely correlated with the PARP1 expression level in the solid tumors^76, 77^. In this regard, it is important to note that PARP1 is highly expressed in SCLC (e.g., compared to non-small cell lung cancer)^77, 78^, suggesting that PARPi (e.g., talazoparib) could potentially be enriched in SCLC tumors. These pharmacodynamic properties of talazoparib (and also other PARP1 inhibitors) in SCLC will be addressed in future studies. Previous studies and our current study show that there is considerable heterogeneity among SCLC in their response to PARPi^64^. However, we found that PARPi was able to induce robust DDR (as shown by γH2AX) in all SCLC cell lines tested, regardless of their sensitivity to PARPi (Fig. 6A; Supplementary Fig. 6D). These observations raise the hypothesis that the selective toxicity of PARPi in certain SCLC does not originate from a cell-specific generation of the DNA damage signal, but rather how each SCLC cell line senses and responds to this genotoxic stimulus. We found that the selective toxicity of PARPi could be explained, at least in part, by the cell-specific regulatory mechanisms of the PiPS proteins. First, among the ASCL1^high^ SCLC cells, ASCL1 was degraded by PARPi in PARPi-sensitive cells, but not in those cells that were resistant to PARPi. Indeed, PARPi-sensitive cells appeared to express higher levels of HUWE1, compared to PARPi-resistant cells. Second, we also found that compared to PARPi-resistant cells, PARPi-sensitive cells expressed higher levels of KDM4A, suggesting that the survival of these cells could be more dependent on the levels of KDM4A. However, how the initial expression of HUWE1 and KDM4A is regulated in SCLC is poorly understood, which will be addressed in future studies. In this context, a recent study showed that SCLC cell lines with high expression of a protein called SLFN11 are sensitive to PARPi^83^. SLFN11 has been shown to interact with RPA1, which leads to the destabilization of RPA1-ssDNA complexes. This results in a “*BRCA*-like” state that renders these cells sensitive to PARPi treatment. In summary, these results suggest that a combination of all these factors might predict the therapeutic response of SCLC to PARPi. Future clinical investigation is warranted to define the full biomarker potential of these protein factors.

In conclusion, using quantitative mass spectrometry, we performed proteomic characterization of the adaptive response of SCLC to PARPi (Fig. 7). Through these unbiased studies, we identified a set of PARPi-responsive proteins that contained key lineage-specific oncoproteins for SCLC (e.g., ASCL1, NEUROD1, POU2F3, KDM4A and KDM5B). Combinatorial treatment of PARPi with the agents targeting the PiPS proteins showed synergistic effects. We then discovered that PARPi induces the degradation of these proteins via a set of DDR-responsive E3 ligases (e.g., HUWE1 and RNF8). Finally, we found that the context-specific sensitivity of SCLC to PARPi originates from the heterogeneous expression of the PiPS proteins (i.e., KDM4A), or their relevant E3 ligases (e.g., HUWE1). Taken together, our efforts highlight PARP1 as a relevant therapeutic target for SCLC. We also show the potential of targeting PiPS proteins and their related UPS machinery in this context. The identification of the molecular underpinnings of these processes could provide novel patient stratification strategies, to accelerate the clinical utility of PARPi in SCLC. Taken together, our data will serve as an invaluable resource, providing the foundation for future hypothesis-driven research that helps delineate the molecular mechanisms that underlie the toxicity of PARPi in SCLC, and more broadly, *BRCA1/2*-proficient cancers.

**Fig. 7.**
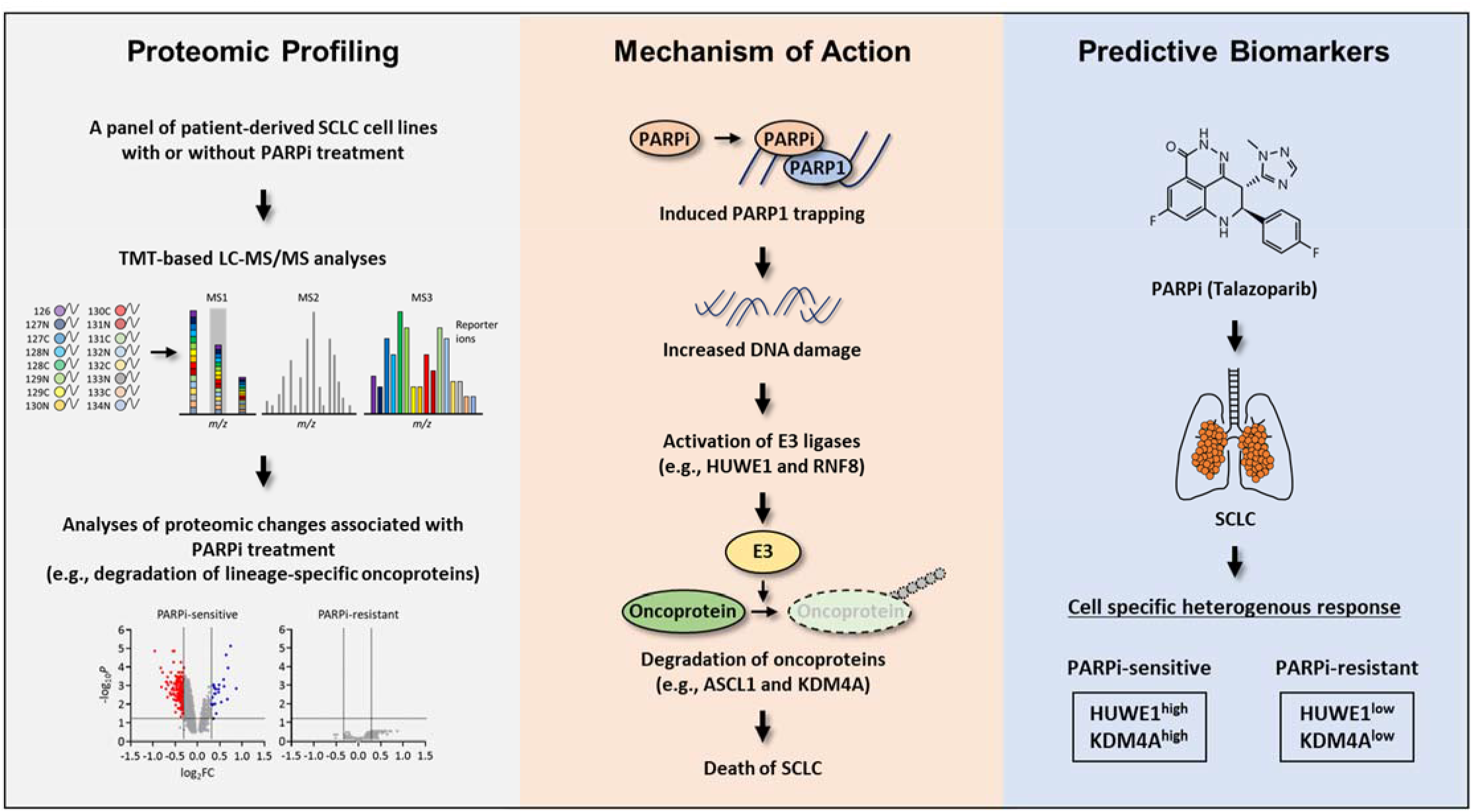
Summary.

## Methods

### Cell lines and culture procedures

Cells used in this study: Human patient-derived small cell lung cancer cell lines NCI-H209, NCI-H1876, NCI-H2107, NCI-H2081, NCI-H128, NCI-H69, NCI-H1092, DMS-79, NCI-H2171, NCI-H524, NCI-H446, NCI-H1048, NCI-H1436, NCI-H889, NCI-H1836, NCI-H2029, SHP-77, NCI-H196, NCI-H1963, NCI-H378, NCI-H82, SW1271, NCI-H1341, and NCI-H841 were either obtained from the Hamon Center for Therapeutic Oncology Research at UT Southwestern medical Center, or purchased from ATCC. Cells were maintained in Roswell Park Memorial Institute (RPMI) 1640 (MilliporeSigma) supplemented with 10% Fetal Bovine Serum (FBS; MilliporeSigma) at 37°C in 5% CO_2_. Human Embryonic Kidney 293T (HEK293T), HEK293TD, and HeLa cell lines were maintained in high glucose Dulbecco’s Modified Eagle’s Medium (DMEM; MilliporeSigma) supplemented with 10% FBS (MilliporeSigma) at 37°C in 5% CO_2_. All cells were found to be mycoplasma-free using the e-Myco kit (Boca Scientific).

### Mice

All animal experiments were performed in compliance with the Institutional Animal Care and Use Committee (IACUC) at the UT Southwestern Medical Center. Female NOD scid gamma (NSG, NOD-SCID) mice (The Jackson Laboratory) at 6-8 weeks of age were used.

### Antibodies and reagents

Antibodies against the following proteins were used (See also Supplementary Table S5). Cell Signaling Technology: PARP1 (#9542), cleaved PARP1 (Asp214, #9541), γH2AX (#9718), cleaved caspase 3 (Asp175, #9661L), α-Tubulin (#3873), Histone H3 (#4499), NEUROD1 (#4373), KDM4A (#5328), KDM5B (#3273), Myc (#2276), HA (#3724), HUWE1 (#5695), MCL-1 (#94296), Tri-Methyl-Histone H3 (Lys9, #13969), Tri-Methyl-Histone H3 (Lys36, #4909), Tri-Methyl-Histone H3 (Lys4, #9751), Phospho-Akt (Ser473, #9271), and Akt (#9272); Santa Cruz Biotechnology: GAPDH (#sc-32233), ASCL1 (#sc-374550), POU2F3 (#sc-293402), UBE2T (#sc-100623), Myc (#sc-60), ubiquitin (#sc-8017), and RNF8 (#sc-271462); Trevigen: PAR (#4335-MC-100); MilliporeSigma: Flag (#F7425); MyBioSource: ZFAND5 (MBS2560210). The following reagents were used (1 μM for 48 hrs, if not indicated): Talazoparib (Pfizer Inc.), BMS-754807, MG132 (10 μM), and Z-VAD-FMK (50 μM) were all purchased from Selleck; Dimethyl sulfoxide and Lipofectamine 2000 were all purchased from Thermo Fisher Scientific; ML324 was purchased from APExBIO; JQ-1 was purchased from AddoQ Bioscience; PBIT, Cycloheximide (10 μg/ml), Polybrene (8 μg/ml), Puromycin (2 μg/ml), NEM (25 mM), sodium orthovanadate (2 mM), and sodium fluoride (20 mM) were purchased from MilliporeSigma.

### Sample preparation for mass spectrometry

All SCLC cell lines were treated with either talazoparib or a panel of small-molecule compounds for 48 hrs. Cells were lysed with 1% SDS lysis buffer containing 10 mM HEPES, pH 7.0, 2 mM MgCl2, 20 U/mL universal nucleases. Protein concentrations were determined with the BCA assay (Thermo Fisher Scientific). Samples were reduced with 3 mM dithiothreitol (DTT) for 20 min and alkylated with 25 mM iodoacetamide (IDA) for 30 min at room temperature (RT) in dark. The detergents were removed by methanol/chloroform precipitation. The proteins were re-solubilized in 8 M urea and digested by Lys-C at a 1:100 (w/w) enzyme/protein ratio for 2 h, followed by trypsin digestion at 1:100 (w/w) enzyme/protein ratio overnight at RT in 2 M urea. The peptides were desalted using Oasis HLB solid-phase extraction (SPE) cartridges (Waters) and approximately 100 μg of peptides for each sample were resuspended in 200 mM HEPES, pH 8.5. The peptides were then labeled with either the amine-based TMT 16-plex reagents (Thermo Fisher Scientific) for 1 hr at RT. Hydroxylamine solution was added to quench the reaction and the labeled peptide samples were combined. Next, the TMT samples were lyophilized and were reconstituted in buffer A (10 mM Ammonium formate, pH 10.0). It was then centrifuged at 10,000 × g for 3 min using Corning® Costar® Spin-X® Plastic Centrifuge Tube Filters (MilliporeSigma) prior to loading onto a ZORBAX 300 Extend-C18 HPLC column (Agilent, Narrow Bore RR 2.1 mm x 100 mm, 3.5 μm particle size, 300 Á pore size). Peptides were fractioned by bRPLC (basic pH reversed-phase HPLC) at a 0.2 mL/min flow rate using a gradient from 0% to 70% buffer B (1% Ammonium formate, pH 10.0 and 90% Acetonitrile). The collected seventeen fractions were lyophilized, desalted, and analyzed by LC-MS/MS as described previously ^84^. Briefly, peptides were separated on a PicoFrit® microcapillary column (75 μm × 15 cm, New Objective). A 180 min linear gradient was developed ranging from 7% to 32% acetonitrile in 0.1% formic acid at 300 nL/min to elute the peptides (Thermo EASY-nLC system).

### Quantitative proteomic analysis by LC-MS/MS

The TMT sample was analyzed by LC-MS/MS on an Orbitrap Eclipse™ Tribrid™ Mass Spectrometer (Thermo Fisher Scientific) using a multi-notch (SPS, synchronous precursor selection)-MS3 approach^85–87^. Briefly, MS1 spectra within 375-1,500 m/z were acquired at 120-K resolving power with a maximum of 50-ms ion injection in the Orbitrap. MS2 spectra were acquired by selection of the most abundant features within 3 sec via collisional induced dissociation in the ion trap using an automatic gain control (AGC) setting of 10-K, quadrupole isolation width of 0.5 m/z and a maximum ion accumulation time of 100 ms. Then an SPS–MS3 scan was performed using up to 10 b- and y-type fragment ions as precursors with an AGC of 200 K for a maximum of 86 ms, with a normalized collision energy setting of 45. MS spectra were searched against a composite database of human protein sequences (Uniprot) and their reversed complement using the Sequest algorithm (Ver28) embedded in an in-house-developed software suite^88^ or Proteome Discoverer™ Software (Thermo Fisher Scientific). MS1 mass tolerance was set to be 50 ppm. Search parameters allowed for full tryptic peptides with a static modification of 57.02146 Da on cystine (Carbamidomethyl), a variable modification of 15.994915 Da on methionine (oxidation), and a static modification of TMT labels (295.1896 Da, TMT 16-plex) on peptide N-terminus and lysine. Search results were filtered to include < 1% matches (both peptide and protein level filtering) to the reverse database by the linear discriminator function using parameters including Xcorr, dCN, missed cleavage, charge state (exclude 1+peptides), mass accuracy, peptide length, and fraction of ions matched to MS/MS spectra. Peptide quantification was performed by using the CoreQuant algorithm implemented in an in-house-developed software suite^89^ or Proteome Discoverer™ Software (Thermo Fisher Scientific). The labeling scheme for the TMT experiments is listed in Supplementary Table S2. For TMT quantification, a 0.03 Th window was scanned around the theoretical m/z of each reporter ion to detect the presence of these ions. The maximum intensity of each ion was extracted, and the signal-to-noise (SN) value of each protein was calculated by summing the reporter ion counts across all identified peptides. Because the same amount of peptides was used for each TMT channel, the total reporter ion intensity of each channel was summed across all quantified proteins, and was then normalized and reported. Data were exported to Excel for further analysis.

### Immunoprecipitation and immunoblot analysis

Cellular lysates were prepared using a Triton X-100 lysis buffer, consisting of 50 mM Tris-HCl, pH 7.4, 150 mM NaCl, 1 mM EDTA, 1 mM N-ethylmaleimide (NEM), 2 mM Na_3_VO_4_, 20 mM NaF, 1 mM PMSF, 1× protease inhibitor cocktail and 0.5% (v/v) Triton X-100. Cellular lysates were clarified by centrifugation at 14,000 × g at 4°C for 15 min. The resulting supernatants were subjected to immunoprecipitation and immunoblot analysis with the corresponding antibodies. For immunoprecipitation, proteins (1 mg) were pre-incubated with protein G Sepharose beads (MilliporeSigma) for pre-clearing and further incubated with 1 – 2 μg of the corresponding antibodies or anti-Flag M2 affinity gel beads (MilliporeSigma) overnight at 4°C. The immunocomplexes were collected with protein G Sepharose beads followed by centrifugation at 3000 × g at 4°C for 2 min. Proteins were eluted from the beads by addition of 2× protein sample buffer, denatured by boiling, separated on SDS-PAGE, and subjected to immunoblot analysis. Indicated antibodies were used. Enhanced chemiluminescence was used to detect specific bands using standard methods. The relative band intensity was measured using the Image J imaging software.

### Cellular fractionation

Cells were fractioned using a subcellular protein fractionation kit (Thermo Fisher Scientific) according to the manufacturer’s instructions. Briefly, cells were harvested with trypsin-EDTA, centrifuged at 500 × g for 5 min, and washed with ice-cold PBS. After adding CEB buffer to the cell pellet, the tube was incubated at 4°C for 10 min with gentle mixing. Following centrifugation at 500 × g for 5 min, the supernatant (cytoplasmic extract) was transferred to a clean pre-chilled tube on ice. Next, the MEB buffer was added to the pellet. The tube was briefly vortexed and was incubated at *4°C* for 10 min with gentle mixing. The tube was then centrifuged at 3000 × g for 5 min and the supernatant (membrane extract) was transferred to a clean pre-chilled tube on ice. An ice-cold NEB buffer was added to the pellet, and the tube was vortexed at the highest setting for 15 s. Following incubation at 4°C for 30 min with gentle mixing, the tube was centrifuged at 5000 × g for 5 min and the supernatant (soluble nuclear extract) was transferred to a clean pre-chilled tube on ice. Lastly, room temperature NEB buffer containing Micrococcal Nuclease and CaCl2 was added to the pellet. The tube was vortexed for 15 s and was incubated at room temperature for 15 min. After incubation, the tube was centrifuged at 16,000 × g for 5 min and the supernatant (chromatin-bound nuclear extract) was transferred to a clean pre-chilled tube on ice.

### *In vivo* drug treatment experiments

All animal experiments were performed in compliance with the Institutional Animal Care and Use Committee (IACUC) at the UT Southwestern Medical Center. Female NOD scid gamma (NSG, NOD-SCID) mice (The Jackson Laboratory) at 6-8 weeks of age were used. Compounds were dissolved in 2% DMSO + 30% PEG 300 + 5% Tween 80 + ddH_2_O (for JQ-1 and ML324) and in 10% DMAc + 6% Solutol + 84% PBS (for talazoparib), respectively. Tumors were engrafted in NSG mice by subcutaneous injection of 1 × 10^6^ cells of H2081 in RPMI1640 medium supplemented with 50% Matrigel (BD Biosciences). Ten days after the injection, mice carrying 100 – 150 mm^3^ subcutaneous tumors were assigned randomly to control and various treatment groups (n = 8–10 for each group). Tumor-bearing mice were orally (for talazoparib) or intraperitoneally injected (for JQ-1 and ML324) with either vehicle, talazoparib (0.3 mg/kg), JQ-1 (25 mg/kg), or ML324 (5 mg/kg) daily for 30 days. The weight of the mice was monitored every 3 days and tumor volume was also measured with calipers every 3 days. Tumor volumes were calculated using a modified ellipsoid formula: Tumor volume = ½ (length × width^2^). Mice were euthanized at 31 days post-injection. Fresh tumor samples were harvested and the weight was measured with an electronic scale, followed by extraction for further experiments.

### Plasmids

Human ASCL1 complementary cDNA was inserted into a pCI/3’Myc (modified pCI vector containing C-terminal Myc tag) by sub-cloning. Flag-Ub and HA-Ub (all in pcDNA3.1), and Flag-HUWE1 WT (Wild-Type) and LD (Ligase-Dead) were kind gifts from Dr. YJ Oh (Yonsei Univ., South Korea), and Dr. Qing Zhong (UT Southwestern Medical Center, USA), respectively. Myc-KDM4A and KDM5B-Myc were kind gifts from Dr. Zhi-Ping Liu (UT Southwestern Medical Center, USA), and Dr. W. Lee Kraus (UT Southwestern Medical Center, USA), respectively. pMH-SFB-RNF8 was purchased from Addgene. Lentiviral plasmids (VSVG and Δ8.9) were kind gifts from A. Kung (Dana Farber Cancer Institute, USA) and D. Baltimore (California Institute of Technology, USA). All plasmids were subjected to DNA sequencing for verification.

### RNA interference and ectopic over-expression using mammalian lentiviral system and transfection

To produce the lentiviruses, shRNA plasmids were co-transfected into HEK293TD cells along with packaging (Δ8.9) and envelope (VSVG) expression plasmids using the Lipofectamine 2000 reagent (Invitrogen) according to the manufacturer’s instructions. After two days, viral supernatants were collected and were filtered using a 0.45-μm filter. Recipient cells were infected in the presence of a serum-containing medium supplemented with 8 μg/ml Polybrene. Two days after infection, cells were used for the indicated experiments. Lipofectamine 2000 reagents were also used to transiently knock-down or over-express the target genes, according to the manufacturer’s instructions. The knock-down or over-expression of target genes was validated by immunoblot assays. The shRNA constructs and over-expression plasmids were listed in Supplementary Table S4 and S5.

### Measurement of IC_50_ (The half maximal inhibitory concentration)

Cells were plated into 96-well plates at densities of 1000 - 2000 cells/well. The next day, cells were treated with the indicated concentration of talazoparib for 4 days. The value of IC_50_ was measured using the CellTiter-Glo assay (Promega) according to the manufacturer’s instructions. Briefly, after incubation, room temperature CellTiter-Glo reagent was added 1:1 to each well, and the plates were incubated at room temperature for 2 min. Luminescence was measured with the Synergy HT Multi-Detection Microplate Reader and was normalized against control cells treated with DMSO.

### Cell viability measurement

Cells were plated into 96-well plates at densities of 1000 - 2000 cells/well. Two days later, cells were treated with indicated compounds. Cell viability was measured using the CellTiter-Glo assay (Promega) according to the manufacturer’s instructions. For synergistic effect measurement, cells were treated with talazoparib, JQ-1, ML324, PBIT, or BMS-754807 alone or in combination as indicated for 1 - 4 days. Briefly, after incubation, room temperature CellTiter-Glo reagent was added 1:1 to each well, and the plates were incubated at room temperature for 2 min. Luminescence was measured with the Synergy HT Multi-Detection Microplate Reader and was normalized against control cells treated with vehicle.

### Quantitative real-time polymerase chain reaction (qRT-PCR)

The mRNA extraction was performed using the RNeasy Mini Kit (QIAGEN) according to the manufacturer’s instructions. Subsequently, total RNAs were converted into cDNA using the SuperScript III Reverse Transcriptase (Thermo Fisher Scientific) following the manual for first-strand cDNA synthesis. qRT-PCR reactions were performed on a CFX384 Touch Real-Time PCR Detection System using 2× Power SYBR Green PCR Master Mix (Thermo Fisher Scientific). For each condition, technical triplicates were prepared and the quantitation cycle (Cq) was calculated. For normalization, GAPDH levels were used as an internal reference and relative expression levels were presented.

### Ubiquitination assays

A cell-based *in vivo* ubiquitination and auto-ubiquitination assays were performed as previously described^90^. In brief, for cell-based *in vivo* ubiquitination assays, cells were incubated with 10 μM MG132 for 6 hrs and were lysed in the abovementioned Triton X-100 lysis buffer. The cell lysates were subjected to immunoprecipitation using the indicated antibodies and beads followed by immunoblot analysis with the corresponding antibodies. For auto-ubiquitination assays, the denaturing IP method was performed. To disrupt non-covalent protein-protein interactions, the cell lysates were heated at 98°C in a lysis buffer containing 1% SDS and then were diluted with Triton X-100 lysis buffer (1:10 ratio).

### DepMap analysis

To determine whether the identified proteins play a role in regulating SCLC cell proliferation and survival, we examined their dependency scores using the data available from the Cancer Dependency Map portal (https://depmap.org/portal/download/)^31^. Gene dependency scores were derived using the DEMETER2 algorithm applied to combined RNAi screen data^91^. A negative gene dependency score corresponds to greater gene essentiality, such that the median gene dependency score of pan-essential genes is normalized to −1 and that of negative control genes is set to 0. We analyzed the distribution of dependency scores from SCLC cell lines (n = 25). Genes were organized in ascending order by median gene dependency scores.

### Drug response data analysis

To quantify the association between PARPi sensitivity and the expression level of certain genes of interest, we performed the analyses focusing on all SCLC cell lines and/or ASCL1^high^ SCLC cell lines which were previously defined as those bearing a significantly higher expression level of the ASCL1 gene^62^. Drug response data across cell lines of small cell lung cancer (SCLC) lineage was download from the Genomics of Drug Sensitivity in Cancer database (GDSC, 61 human SCLC cell lines, https://www.cancerrxgene.org/)^61^. The GDSC cohorts used the half maximal inhibitory concentration (IC_50_) as a measure of the potency of a small molecule compound in inhibiting the growth of a specific cell line.

### Quantification and statistical analysis

All the other statistical analyses including unpaired Student’s t-tests, one- and two-way ANOVA were performed using the GraphPad Prism software (v8.4.2). Data were calculated as mean ± SEM or SD. The following indications of significance were used throughout the manuscript: \**P* < 0.05, \*\**P* < 0.01, \*\*\**P* < 0.001, \*\*\*\**P* < 0.0001, N.S, not significant. For the proteomic data analyses, we first grouped the SCLC cell lines into four subtypes (ASCL1^high^, NEUROD1^high^, POU2F3^high^ and TF^low^). For the ASCL1^high^ and NEUROD1^high^ subtypes, we then divided the cells into two categories based on their sensitivities to talazoparib treatment (i.e., talazoparib-sensitive and talazoparib-resistant). The changes of the protein abundances between DMSO and talazoparib were analyzed in each sub-group using the Limma package (3.50.0) in R (4.1.1). The POU2F3^high^ and TF^low^ subtypes were not further divided because they only contained either talazoparib-sensitive cell lines (POU2F3^high^) or talazoparib-resistant cell lines (TF^low^). The paired t-test analyses were used when comparing PARPi vs DMSO in each group. The significance was calculated, and the resulting *P* values were corrected for multiple testing using the Benjamini–Hochberg method in R.

## Contact for reagent and resource sharing

### Lead contact

Information regarding the request of materials and reagents should be directed to and will be fulfilled by the lead contact and co-responding author, Yonghao Yu.

### Materials availability

Plasmids generated in this study are available upon request. Please email the lead contact and corresponding author, Yonghao Yu.

### Data availability

The mass spectrometry proteomics data have been deposited to the ProteomeXchange Consortium via the PRIDE^64^ partner repository with the dataset identifier PXD029968 (Reviewer account details: Username; Password: IJQrAwNO) and PXD029972 (Reviewer account details:Username; Password: Ra7eAIM8).

## Acknowledgements

We thank Dr. Sherry Niessen for providing talazoparib, Dr. Lee Kraus for providing the KDM5B-Myc plasmid, Dr. Zhi-Ping Liu for providing the Myc-KDM4A plasmid, Dr. Rongkuan Hu for the initiation of the quantitative mass spectrometry experiments, and Dr. Y.J. Oh for providing the Flag- and HA-Ub plasmids. We thank Drs. Sherry Niessen, Shubha Bagrodia, Xia Ding, and Michael White (all current or previous employees of Pfizer) for helpful discussions and for funding this work. This work was supported, in part, by grants from NIH (R21CA261018, R01NS122533, and R35GM134883 to Y.Y. and U01CA213338 to J.M.), Welch foundation (I-1800 to Y.Y.) and a sponsored research agreement from Pfizer.

## Author contributions

C.K. and Y.Y conceived of the research, designed all experiments, and wrote the paper. C.K. performed all experiments including cell biology, biochemical methods, and *in vivo* experiments. C.K., S.W., P.L., Z.Z., and Q.D. performed TMT-MS experiments. J.H. and J.M. provided cell lines. C.K., X.W., Y.L, S.J., and J.K. developed methods and analyzed mass spectrometry data using bioinformatics tools. C.K., H.L., J.M., and Y.Y discussed the work.

## Supplementary information

Supplementary Fig. 1-7

Supplementary Tables S1-S5

## Supplementary tables and data

Supplementary Table S1. The information of patient-derived SCLC cell lines used in this study

Supplementary Table S2. The dataset of the TMT experiments

Supplementary Table S3. The list of PiPS proteins

Supplementary Table S4. Primer sequence

Supplementary Table S5. Key resources table

## Supplementary figure legends

**Supplementary Fig. 1.**
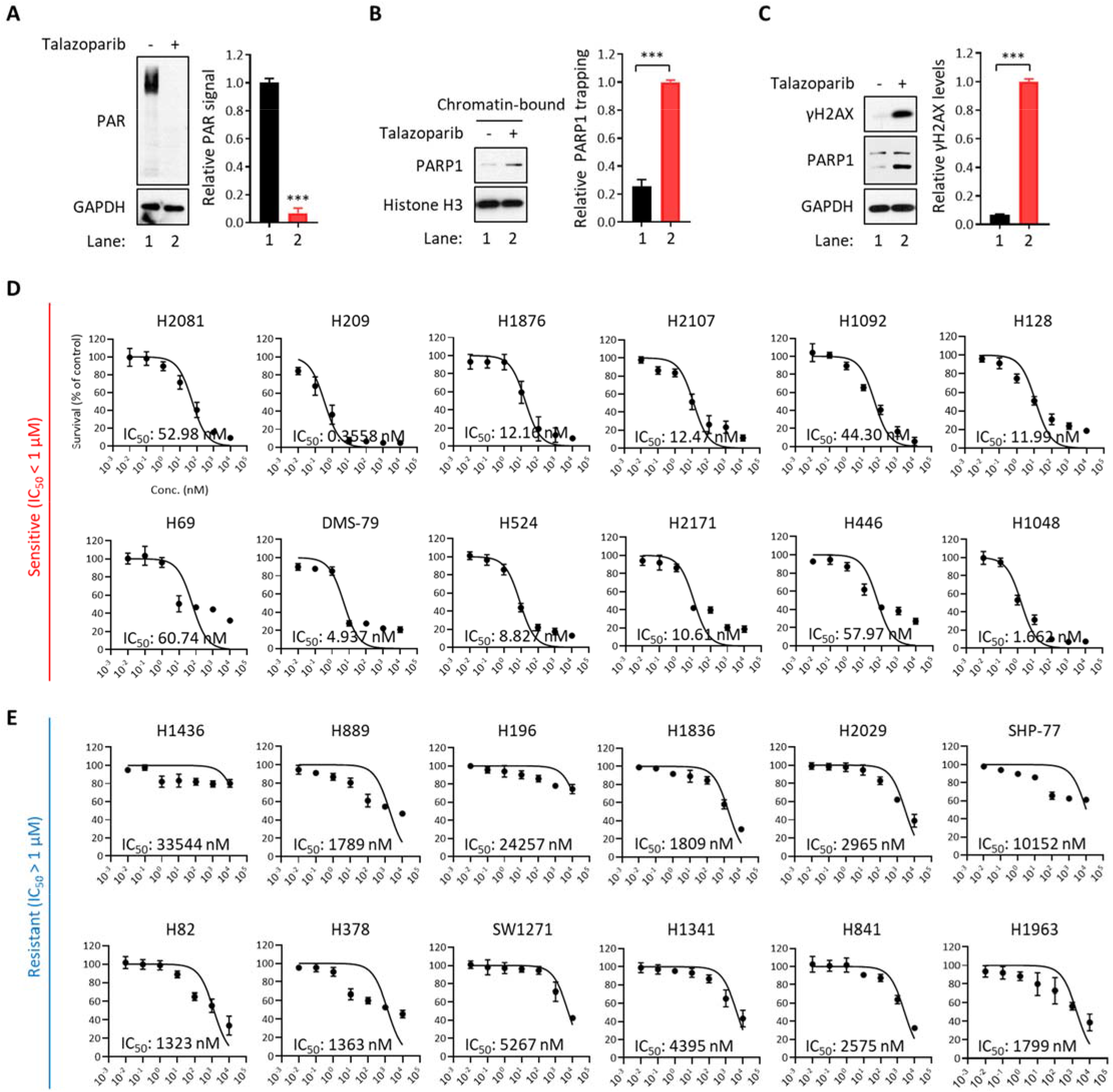
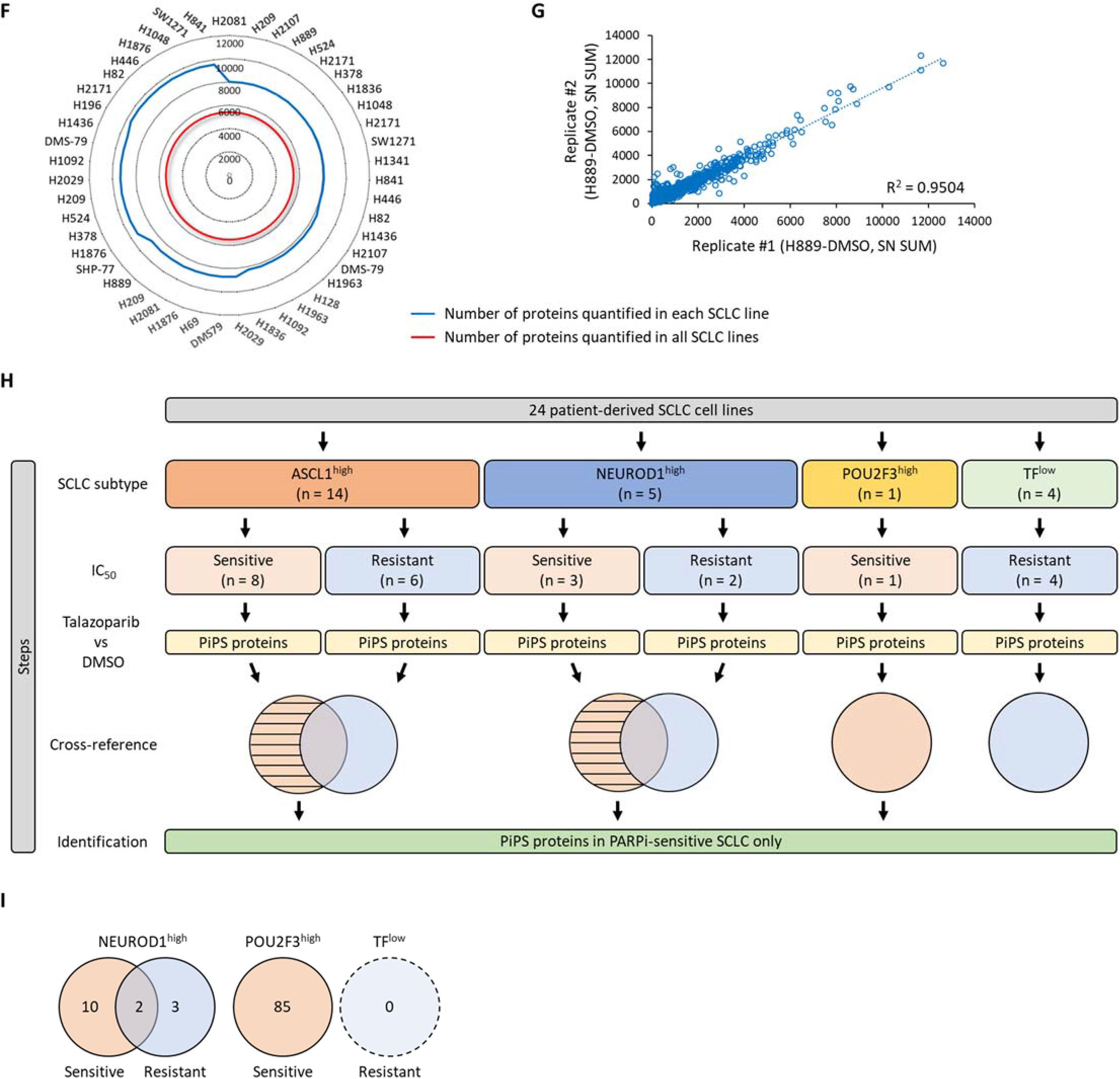
Related to Fig. 1. (A) PAR levels in talazoparib treatment. H2081 cells were treated with talazoparib (1 μM for 48 hrs) and the cell lysates were subjected to immunoblot analysis using the indicated antibodies (n = 3). (B) The levels of PARP1 trapping in talazoparib treatment. H2081 cells were treated with or without talazoparib (1 μM for 48 hrs). Chromatin-bound fractions were isolated and subjected to immunoblot analysis using the indicated antibodies (n = 3). (C) The extent of DNA damage and PARP1 cleavage in talazoparib treatment. H2081 cells were treated with talazoparib (1 μM for 48 hrs) and cell lysates were subjected to immunoblot analysis using the indicated antibodies (n = 3). (D) The concentration-response curves and IC_50_ values in PARPi-sensitive SCLC cell lines (IC_50_ < 1 μM). (E) The concentration-response curves and IC_50_ values in PARPi-resistant SCLC cell lines (IC_50_ > 1 μM). (F) Radar chart showing the number of proteins quantified in this study. See also Supplementary Table S2. (G) Reproducibility of biological replicates in H889 cells. (H) Steps of the identification of PiPS proteins in PARPi-sensitive and -resistant SCLC cell lines of each SCLC subtype. TF^low^, low expression of all three transcription factors. See also Fig. 1C and Supplementary Fig. 1I. (I) PiPS proteins identified in PARPi-sensitive and -resistant SCLC of NEUROD1^high^, POU2F3^high^, and TF^low^ subtype. See also Supplementary Table S3.

**Supplementary Fig. 2.**
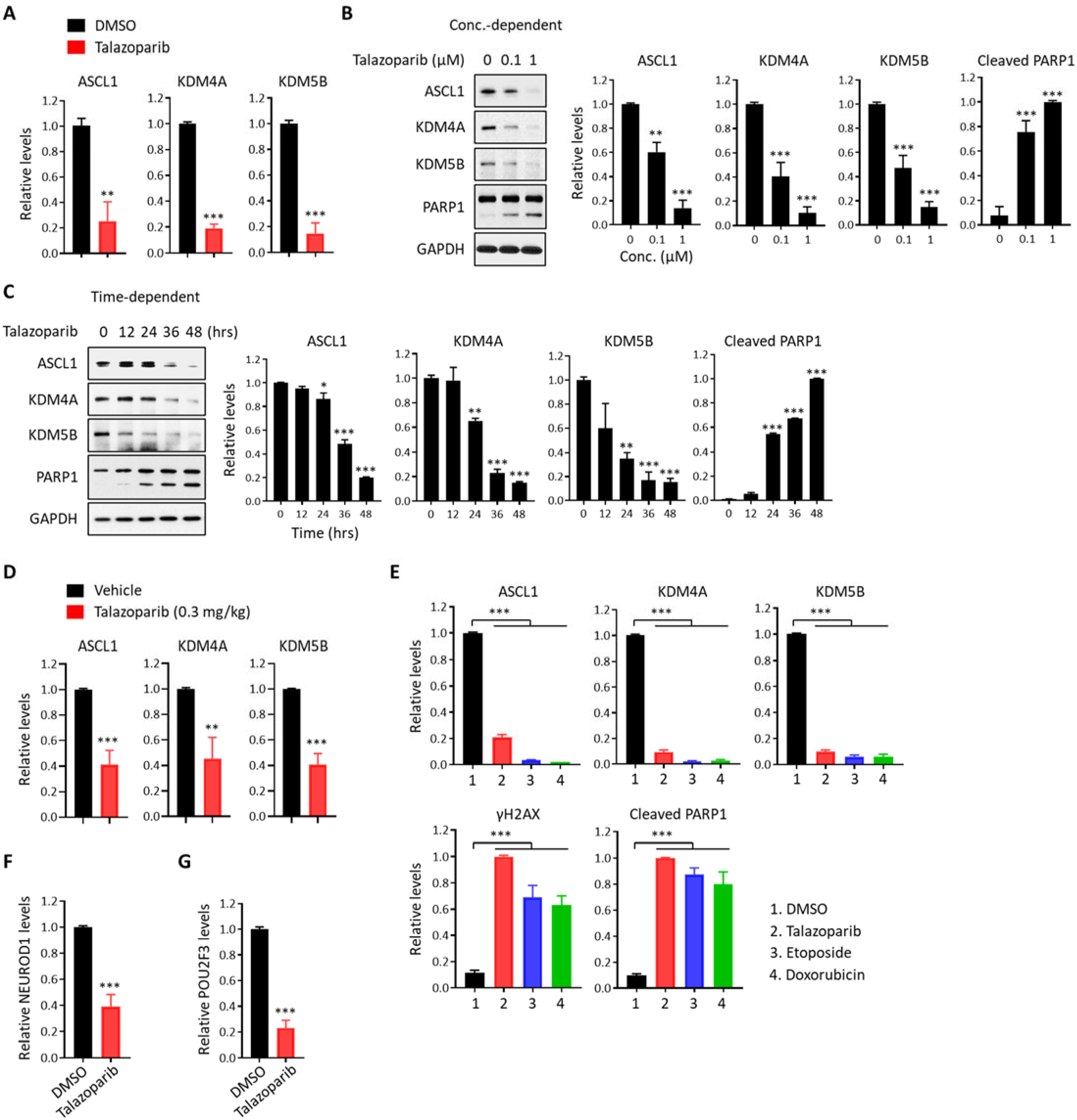
Related to Fig. 2. (A) The levels of ASCL1, KDM4A, and KDM5B in talazoparib treatment. Quantification of the immunoblots presented in Fig. 2A was shown (n = 3). (B) The levels of ASCL1, KDM4A, and KDM5B in a time-dependent manner. H2081 cells were treated with talazoparib (1 μM) in a time-dependent manner and the cell lysates were subjected to immunoblot analysis using the indicated antibodies (n = 3). (C) The levels of ASCL1, KDM4A, and KDM5B in a concentration-dependent manner. H2081 cells were treated with talazoparib (for 48 hrs) in a concentration-dependent manner and the cell lysates were subjected to immunoblot analysis using the indicated antibodies (n = 3). (D) The levels of ASCL1, KDM4A, and KDM5B *in vivo*. Quantification of the immunoblots presented in Fig. 2B was shown (n = 3). (E) The levels of ASCL1, KDM4A, and KDM5B in the treatment of DNA damaging agents. Quantification of the immunoblots presented in Fig. 2E was shown (n = 3). (F) The levels of NEUROD1 in talazoparib treatment. Quantification of the immunoblots presented in Fig. 2F was shown (n = 3). (G) The levels of POU2F3 in talazoparib treatment. Quantification of the immunoblots presented in Fig. 2G was shown (n = 3).

**Supplementary Fig. 3.**
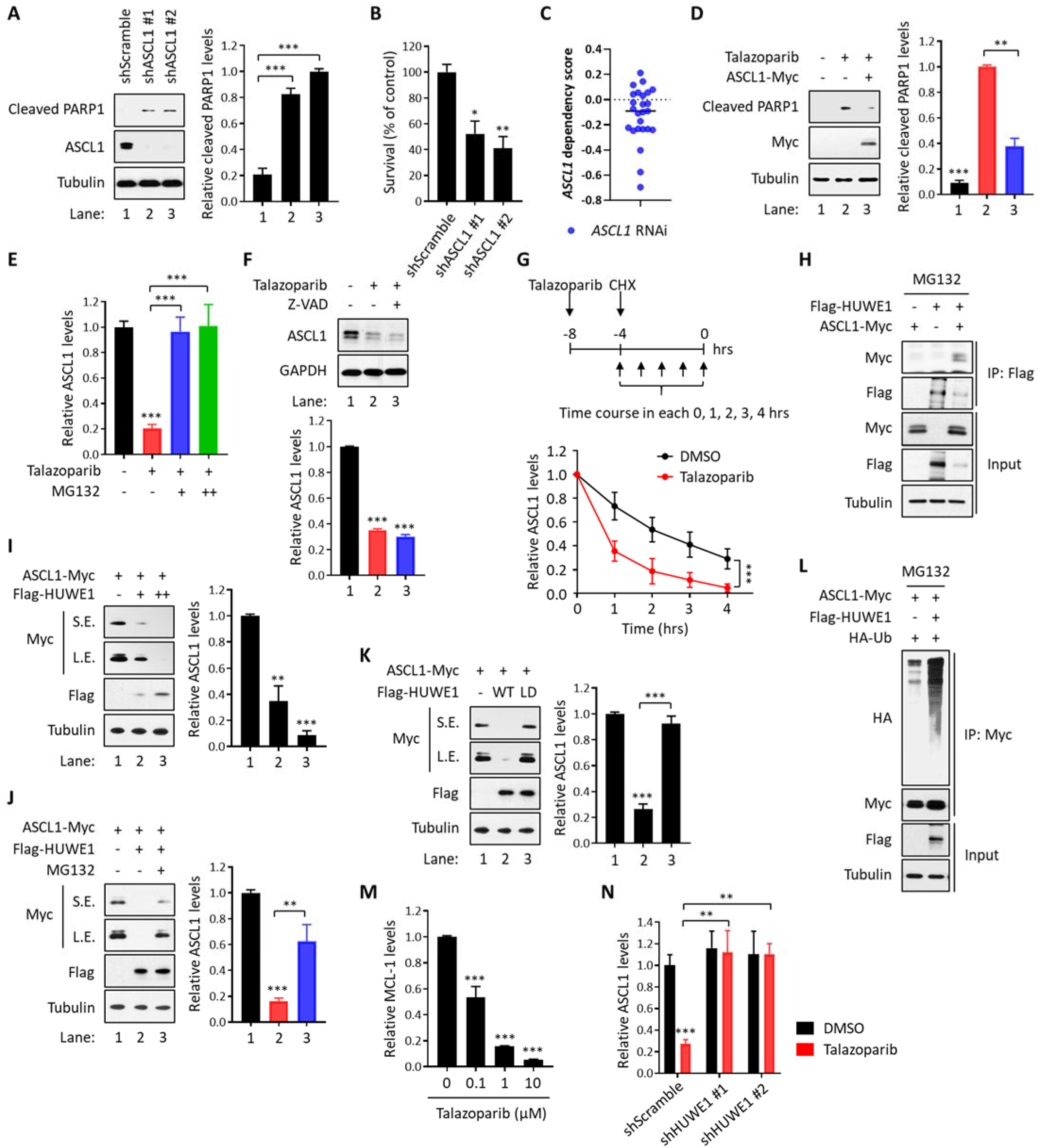
Related to Fig. 3. (A) The levels of cleaved PARP1 in depletion of ASCL1. H2081 cells were depleted with ASCL1 (shASCL1 #1 or #2) and the cell lysates were subjected to immunoblot analysis using the indicated antibodies (n = 3). (B) Viability assays in depletion of ASCL1. H2081 cells were depleted with ASCL1 (shASCL1 #1 or #2) and viability was measured using a CellTiter-Glo assay (n = 3). (C) DepMap analysis for the requirement of ASCL1 in human SCLC cell lines. (D) The levels of cleaved PARP1 in ectopic expression of ASCL1. H2081 cells expressing ASCL1 (ASCL1-Myc) were treated with talazoparib (1 μM for 48 hrs) and the cell lysates were subjected to immunoblot analysis using the indicated antibodies (n = 3). (E) Inhibition of ASCL1 degradation. Quantification of the immunoblots presented in Fig. 3C was shown (n = 3). (F) The levels of ASCL1 in the inhibition of caspase activity. H2081 cells pre-treated with Z-VAD (50 μM for 1 hr) were treated with talazoparib (1 μM for 48 hrs) and the cell lysates were subjected to immunoblot analysis using the indicated antibodies (n = 3). (G) Half-life of ASCL1. Quantification of the immunoblots presented in Fig. 3E was shown (n = 3). Top, schematic flow; Bottom, the normalized levels of ASCL1. (H) The interaction of ASCL1 with HUWE1. HEK293T cells transfected with Flag-HUWE1 or ASCL1-Myc alone or in combination were treated with MG132 (10 μM for 6hr) and the cell lysates were subjected to co-immunoprecipitation using anti-Flag M2 affinity gel beads. Immunoprecipitates or inputs were resolved by SDS-PAGE and subjected to immunoblot analysis using the indicated antibodies. (I) The levels of ASCL1 in ectopic expression of HUWE1. HEK293T cells were transfected with ASCL1-Myc alone or in combination with Flag-HUWE1 and the cell lysates were subjected to immunoblot analysis using the indicated antibodies (n = 3). S.E., short exposure; L.E., long exposure. (J) The levels of ASCL1 with HUWE1 in proteasome inhibition. HEK293T cells transfected with ASCL1-Myc alone or in combination with Flag-HUWE1 were treated with MG132 (10 μM for 6hr). The cell lysates were subjected to immunoblot analysis using the indicated antibodies (n = 3). S.E., short exposure; L.E., long exposure. (K) The levels of ASCL1 in HUWE1 E3 ligase activity. HEK293T cells transfected with ASCL1-Myc alone or in combination with Flag-HUWE1 wild-type (WT) or ligase-dead (LD) mutant were subjected to immunoblot analysis using the indicated antibodies (n = 3). S.E., short exposure; L.E., long exposure. (L) Ubiquitination of ASCL1 by HUWE1. HEK293T cells transfected with ASCL1-Myc alone or in combination with Flag-HUWE1 were treated with MG132 (10 μM for 6hr) and the cell lysates were subjected to immunoprecipitation using anti-Myc antibody. Immunoprecipitates or inputs were resolved by SDS-PAGE and subjected to immunoblot analysis using the indicated antibodies (n = 3). (M) Downregulation of MCL-1 in talazoparib treatment. Quantification of the immunoblots presented in Fig. 3H was shown (n = 3). (N) Inhibition of ASCL1 degradation. Quantification of the immunoblots presented in Fig. 3J was shown (n = 3).

**Supplementary Fig. 4.**
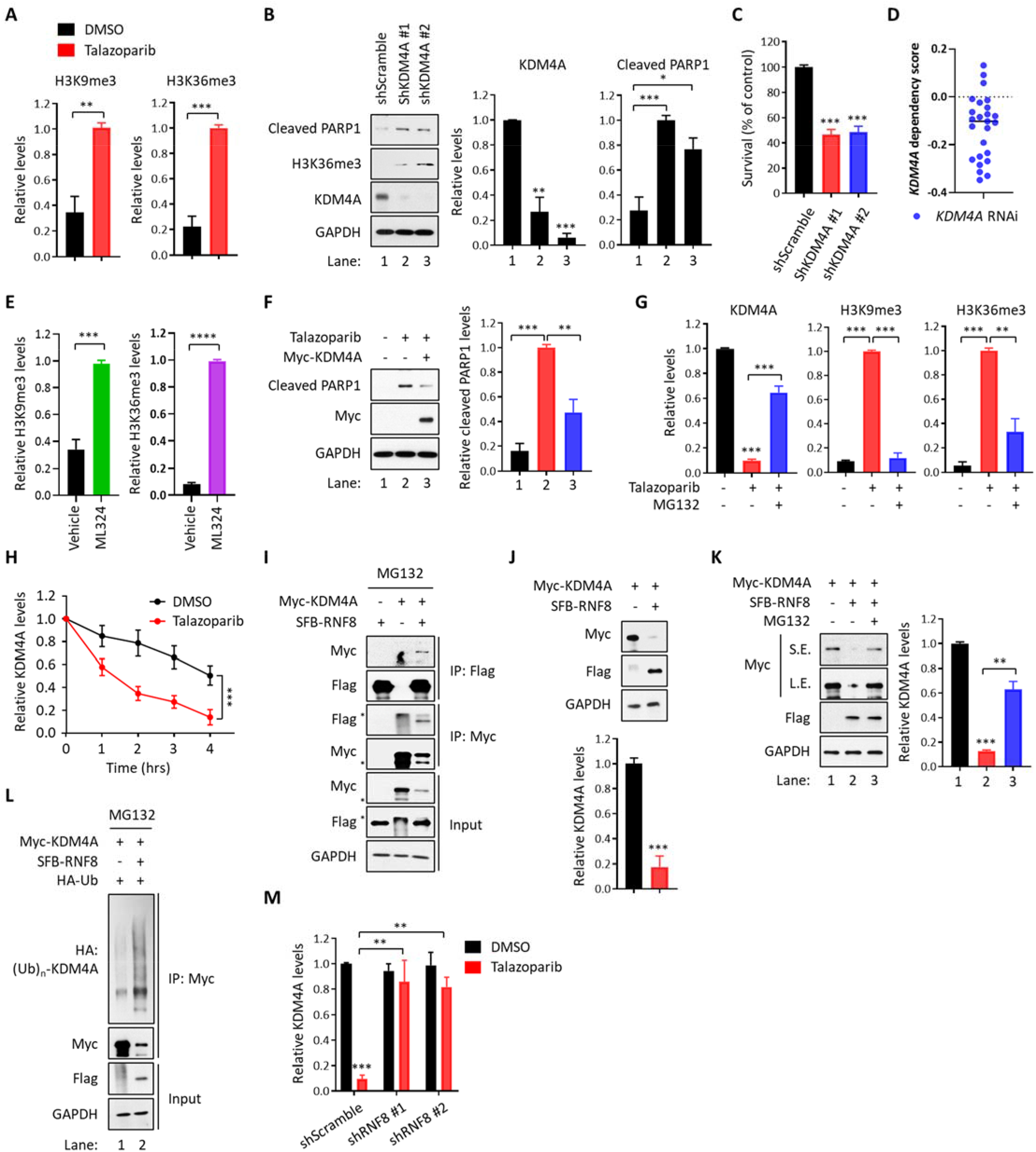
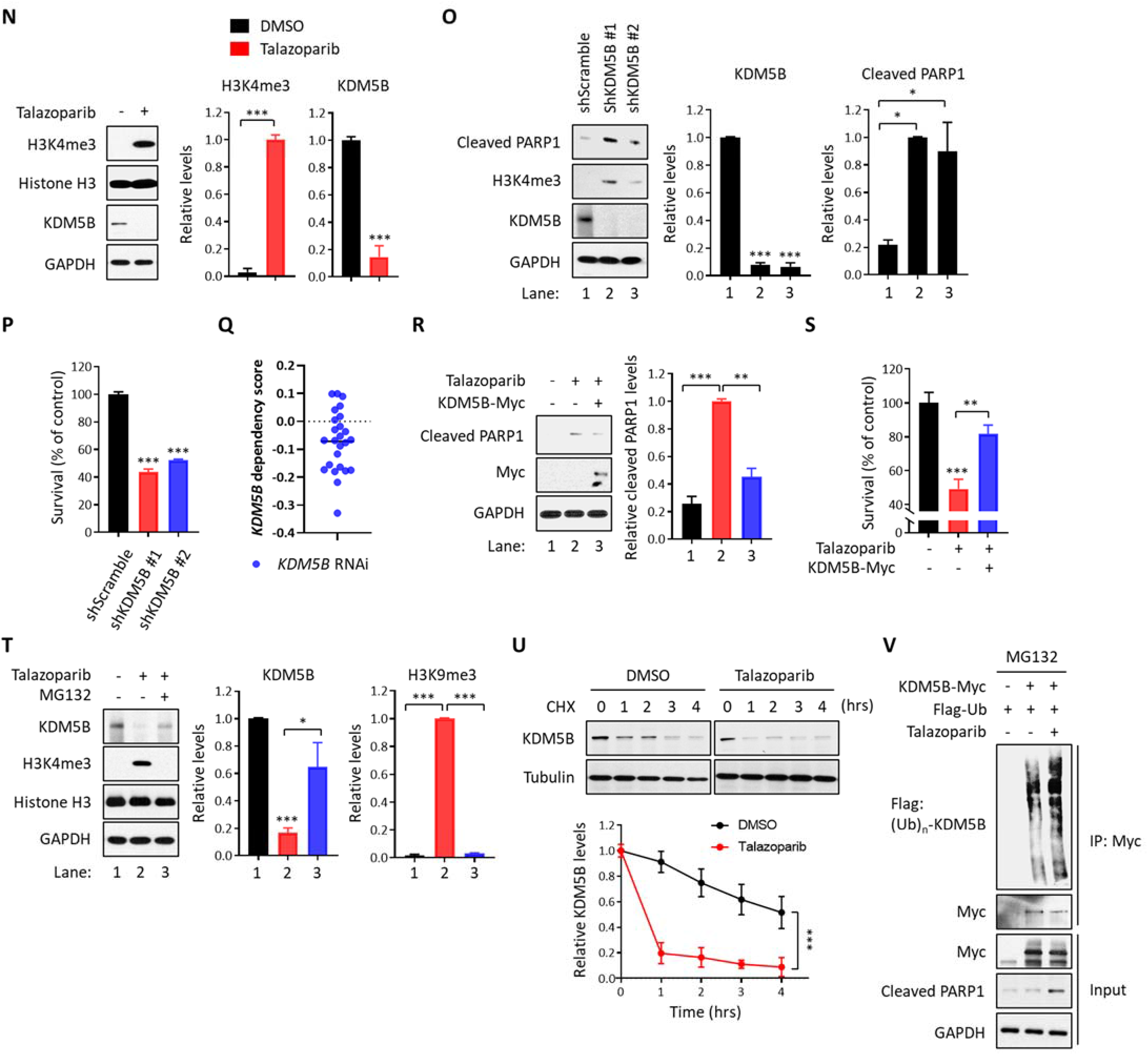
Related to Fig. 4. (A) The levels of histone trimethylation (H3K9me3 and H3K36me3) in talazoparib treatment. Quantification of the immunoblots presented in Fig. 4A was shown (n = 3). (B) The levels of cleaved PARP1 and Histone H3 trimethylation (H3K36me3) in depletion of KDM4A. H2081 cells were depleted with KDM4A (shKDM4A #1 or #2) and the cell lysates were subjected to immunoblot analysis using the indicated antibodies (n = 3). (C) Viability assays in depletion of KDM4A. H2081 cells were depleted with KDM4A (shKDM4A #1 or #2) and viability was measured using a CellTiter-Glo assay (n = 3). (D) DepMap analysis for the requirement of KDM4A in human SCLC cell lines. (E) The levels of histone trimethylation (H3K9me3 and H3K36me3) *in vivo*. Quantification of the immunoblots presented in Fig. 4C was shown (n = 3). (F) The levels of cleaved PARP1 in ectopic expression of KDM4A. H2081 cells expressing KDM4A (Myc-KDM4A) were treated with talazoparib (1 μM for 48 hrs) and the cell lysates were subjected to immunoblot analysis using the indicated antibodies (n = 3). (G) The levels of KDM4A degradation and its subsequent histone trimethylation in proteasome inhibition. Quantification of the immunoblots presented in Fig. 4F was shown (n = 3). (H) The half-life of KDM4A. Quantification of the immunoblots presented in Fig. 4G was shown (n = 3). (I) The interaction of KDM4A with RNF8. HEK293T cells transfected with Myc-KDM4A or SFB-RNF8 alone or in combination were treated with MG132 (10 μM for 6hr) and the cell lysates were subjected to co-immunoprecipitation using anti-Myc antibody and anti-Flag M2 affinity gel beads. Immunoprecipitates or inputs were resolved by SDS-PAGE and subjected to immunoblot analysis using the indicated antibodies. The asterisks indicate non-specific bands. (J) The levels of KDM4A in ectopic expression of RNF8. HEK293T cells were transfected with Myc-KDM4A alone or in combination with SFB-RNF8 and the cell lysates were subjected to immunoblot analysis using the indicated antibodies (n = 3). (K) The levels of KDM4A with RNF8 in proteasome inhibition. HEK293T cells transfected with Myc-KDM4A alone or in combination with SFB-RNF8 were treated with MG132 (10 μM for 6hr). The cell lysates were subjected to immunoblot analysis using the indicated antibodies (n = 3). S.E., short exposure; L.E., long exposure. (L) Ubiquitination of KDM4A by RNF8. HEK293T cells transfected with Myc-KDM4A alone or in combination with SFB-RNF8 were treated with MG132 (10 μM for 6hr) and the cell lysates were subjected to immunoprecipitation using anti-Myc antibody. Immunoprecipitates or inputs were resolved by SDS-PAGE and subjected to immunoblot analysis using the indicated antibodies (n = 3). (M) Inhibition of KDM4A degradation. Quantification of the immunoblots presented in Fig. 4K was shown (n = 3). (N) The levels of KDM5B and histone H3 trimethylation (H3K4me3) in talazoparib treatment. H2081 cells were treated with or without talazoparib (1 μM for 48 hrs) and the cell lysates were subjected to immunoblot analysis using the indicated antibodies (n = 3). (O) The levels of cleaved PARP1 and histone H3 trimethylation (H3K4me3) in depletion of KDM5B. H2081 cells were depleted with KDM5B (shKDM5B #1 or #2) and the cell lysates were subjected to immunoblot analysis using the indicated antibodies (n = 3). (P) Viability assays in depletion of KDM5B. H2081 cells were depleted with KDM5B (shKDM5B #1 or #2) and viability was measured using a CellTiter-Glo assay (n = 3). (Q) DepMap analysis for the requirement of KDM5B in human SCLC cell lines. (R) The levels of cleaved PARP1 in ectopic expression of KDM5B. H2081 cells expressing KDM5B (KDM5B-Myc) were treated with talazoparib (1 μM for 48 hrs) and the cell lysates were subjected to immunoblot analysis using the indicated antibodies (n = 3). (S) Viability assays in ectopic expression of KDM5B. H2081 cells expressing KDM5B (KDM5B-Myc) were treated with talazoparib (1 μM for 48 hrs) and viability was measured using a CellTiter-Glo assay (n = 3). (T) The levels of KDM5B and histone H3 trimethylation (H3K9me3) in proteasome inhibition. H2081 cells were treated with talazoparib (1 μM for 48 hrs) and then, MG132 (10 μM) was further treated for last 6 hrs. The cell lysates were subjected to immunoblot analysis using the indicated antibodies (n = 3). (U) The half-life of KDM5B. H2081 cells pre-treated with or without talazoparib (1 μM) were treated with CHX (10 μg/ml) as indicated. The cell lysates were subjected to immunoblot analysis using the indicated antibodies (n = 3). (V) Ubiquitination of KDM5B in talazoparib treatment. H2081 cells expressing KDM5B (KDM5B-Myc) were treated with talazoparib (1 μM for 48 hrs) and then, proteasome inhibitor (MG132, 10 μM) was further treated for last 6 hrs. The cell lysates were subjected to immunoprecipitation analysis using anti-Myc antibody. Immunoprecipitates or inputs were resolved by SDS-PAGE and subjected to immunoblot analysis using the indicated antibodies.

**Supplementary Fig. 5.**
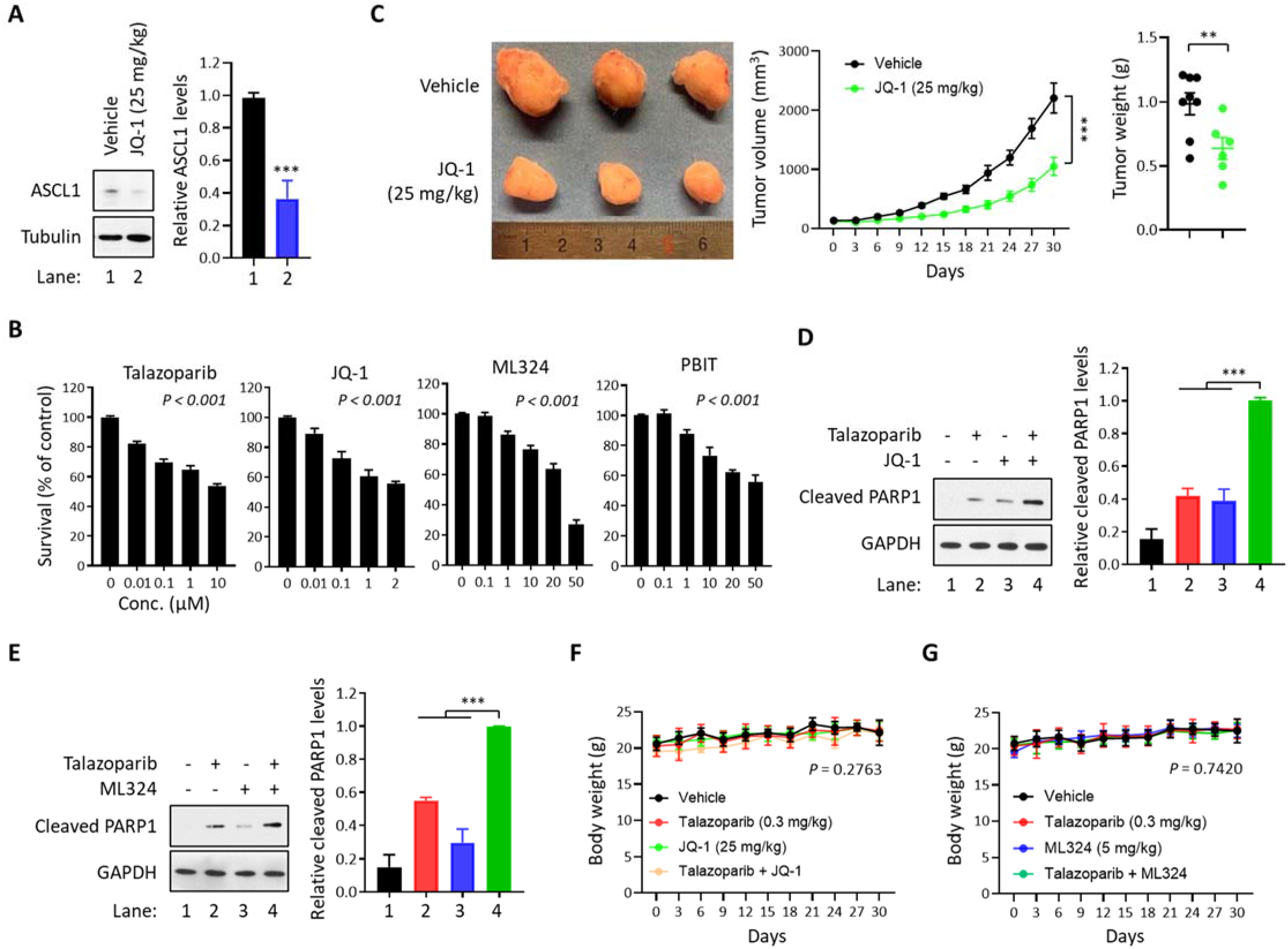
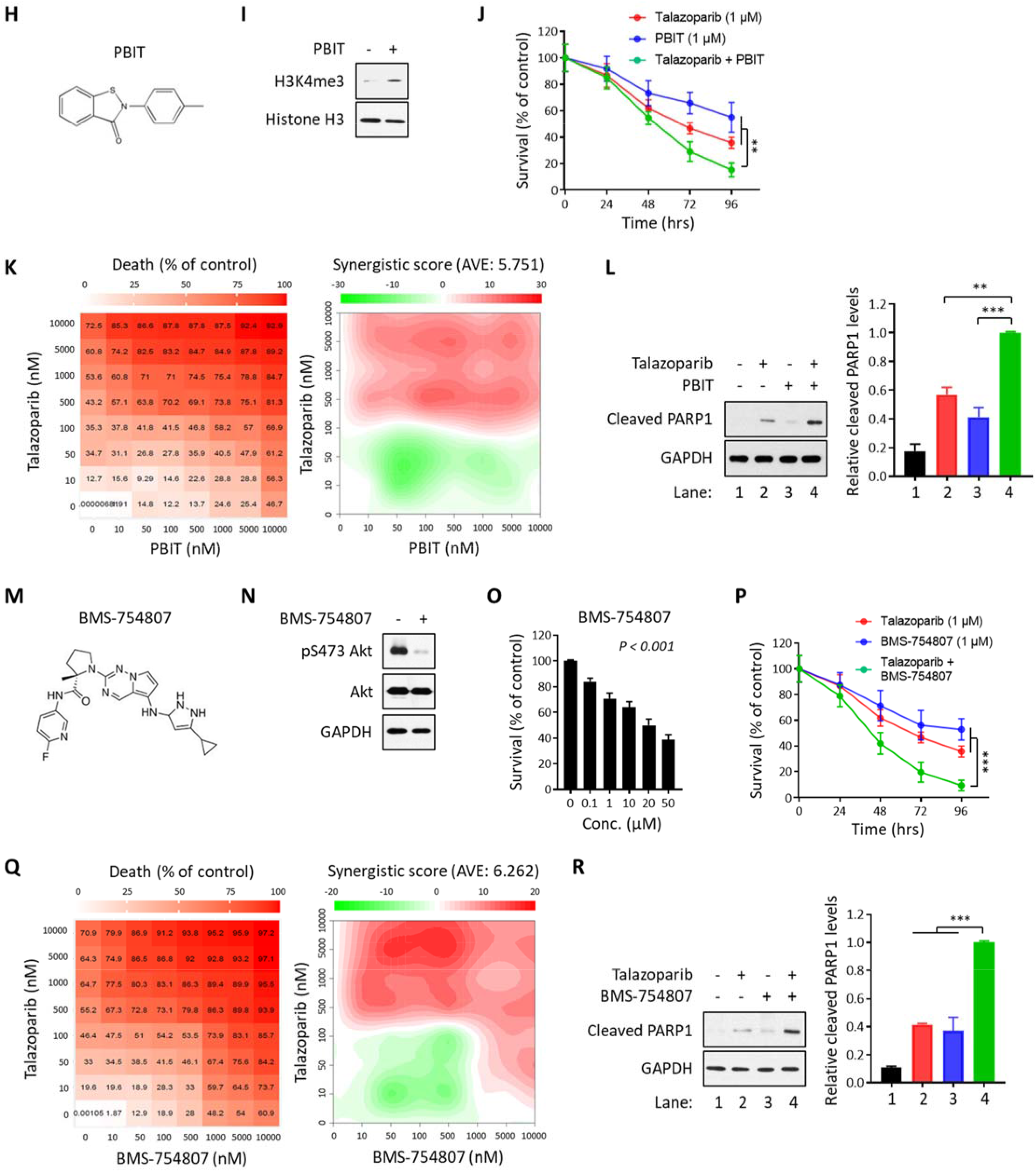
Related to Fig. 5. (A) The levels of ASCL1 *in vivo*. H2081-implanted xenograft tumors were treated with or without JQ-1 (25 mg/kg for 30 days) and the tumor extracts were subjected to immunoblot analysis using the indicated antibodies (n = 8). (B) The cytotoxic effects of talazoparib, JQ-1, ML324, or PBIT in a concentration-dependent manner. H2081 cells were treated with talazoparib, JQ-1, ML324, or PBIT for 48 hrs in a concentration-dependent manner and viability was measured using a CellTiter-Glo assay (n = 3). (C) Toxicity of JQ-1 *in vivo*. Mice implanted with H2081 cells were treated with or without JQ-1 (25 mg/kg for 30 days). Left, the image of tumor size; Right, tumor volume and weight (n = 8–10 per each group). (D) The synergistic effect of talazoparib with JQ-1. H2081 cells treated with talazoparib (1 μM for 48 hrs), JQ-1 (0.1 μM for 48 hrs), or talazoparib + JQ-1 were subjected to immunoblot analysis using the indicated antibodies (n = 3). (E) The synergistic effect of talazoparib with ML324. H2081 cells treated with talazoparib (1 μM for 48 hrs), ML-324 (1 μM for 48 hrs), or talazoparib + ML324 were subjected to immunoblot analysis using the indicated antibodies (n = 3). (F) Body weight of H2081-implanted xenograft tumors to talazoparib (0.3 mg/kg for 30 days), JQ-1 (25 mg/kg for 30 days), and talazoparib + JQ-1. (G) Body weight of H2081-implanted xenograft tumors to talazoparib (0.3 mg/kg for 30 days), ML324 (5 mg/kg for 30 days), and talazoparib + ML324. (H) The structure of PBIT. (I) The levels of histone H3 trimethylation (H3K4me3) in PBIT treatment. H2081 cells were treated with PBIT (1 μM for 48 hrs) and the cell lysates were subjected to immunoblot analysis using the indicated antibodies. (J) The synergistic effect of talazoparib with PBIT in a time-dependent manner. H2081 cells treated with talazoparib (1 μM), PBIT (1 μM), or talazoparib + PBIT in a time-dependent manner. Viability was measured using a CellTiter-Glo assay (n = 3). (K) The synergistic effect of talazoparib with PBIT in a concentration-dependent manner. H2081 cells treated with talazoparib, PBIT, or talazoparib + PBIT in a concentration-dependent manner. Viability was measured using a CellTiter-Glo assay (n = 3). Left, cell death ratio; Right, synergistic score. (L) The synergistic effect of talazoparib with PBIT. H2081 cells treated with talazoparib (1 μM for 48 hrs), PBIT (1 μM for 48 hrs), or talazoparib + PBIT were subjected to immunoblot analysis using the indicated antibodies (n = 3). (M) The structure of BMS-754807. (N) The levels of pS473 Akt in BMS-754807 treatment. H2081 cells were treated with BMS-754807 (1 μM for 48 hrs) and the cell lysates were subjected to immunoblot analysis using the indicated antibodies. (O) The cytotoxic effects of BMS-754807 in a concentration-dependent manner. H2081 cells were treated with BMS-754807 for 48 hrs in a concentration-dependent manner and viability was measured using a CellTiter-Glo assay (n = 3). (P) The synergistic effect of talazoparib with BMS-754807 in a time-dependent manner. H2081 cells treated with talazoparib (1 μM), BMS-754807 (1 μM), or talazoparib + BMS-754807 in a time-dependent manner. Viability was measured using a CellTiter-Glo assay (n = 3). (Q) The synergistic effect of talazoparib with BMS-754807 in a concentration-dependent manner. H2081 cells treated with talazoparib, BMS-754807, or talazoparib + BMS-754807 in a concentration-dependent manner. Viability was measured using a CellTiter-Glo assay (n = 3). Left, cell death ratio; Right, synergistic score. (R) The synergistic effect of talazoparib with BMS-754807. H2081 cells treated with talazoparib (1 μM for 48 hrs), BMS-754807 (1 μM for 48 hrs), or talazoparib + BMS-754807 were subjected to immunoblot analysis using the indicated antibodies (n = 3).

**Supplementary Fig. 6.**
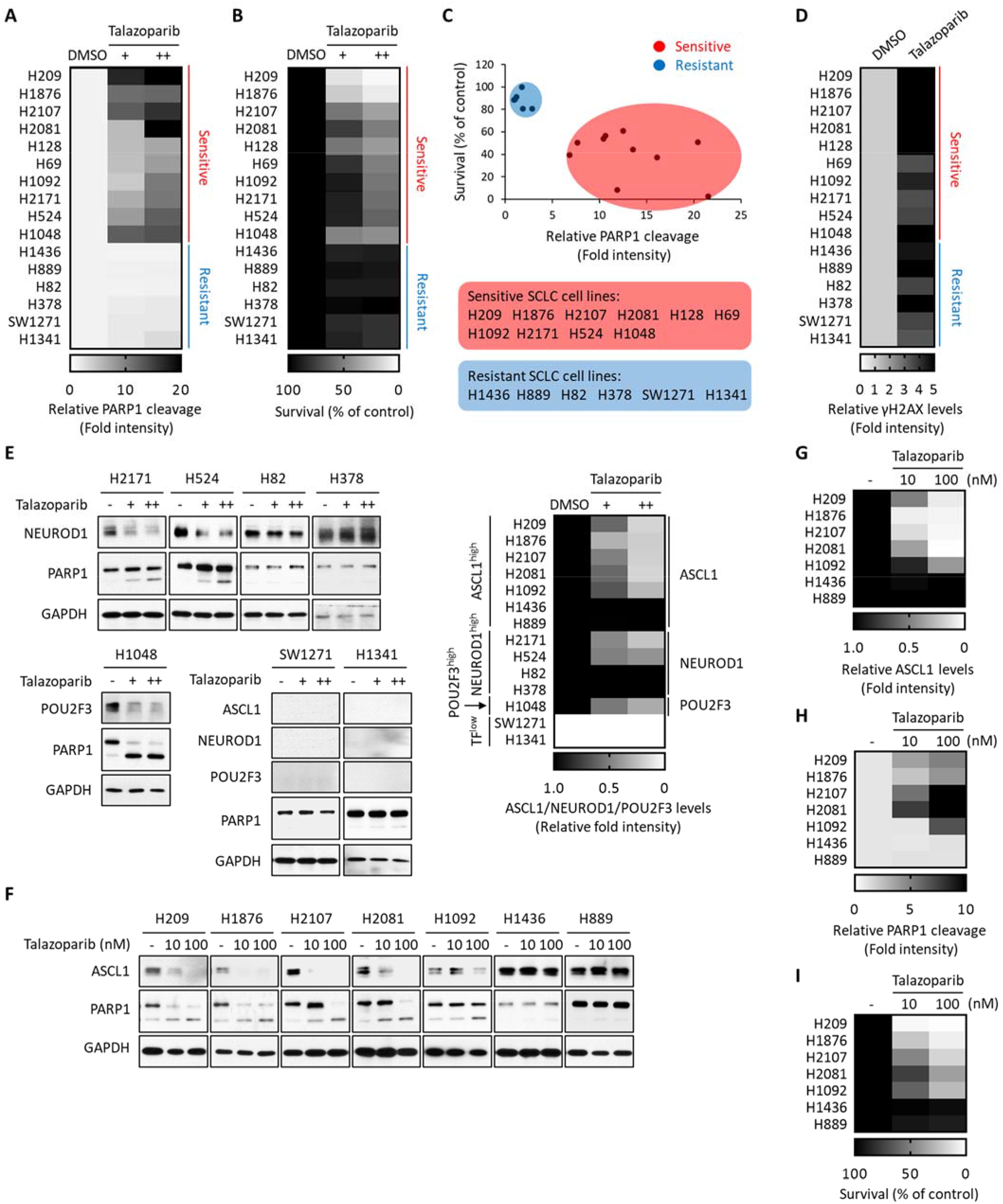
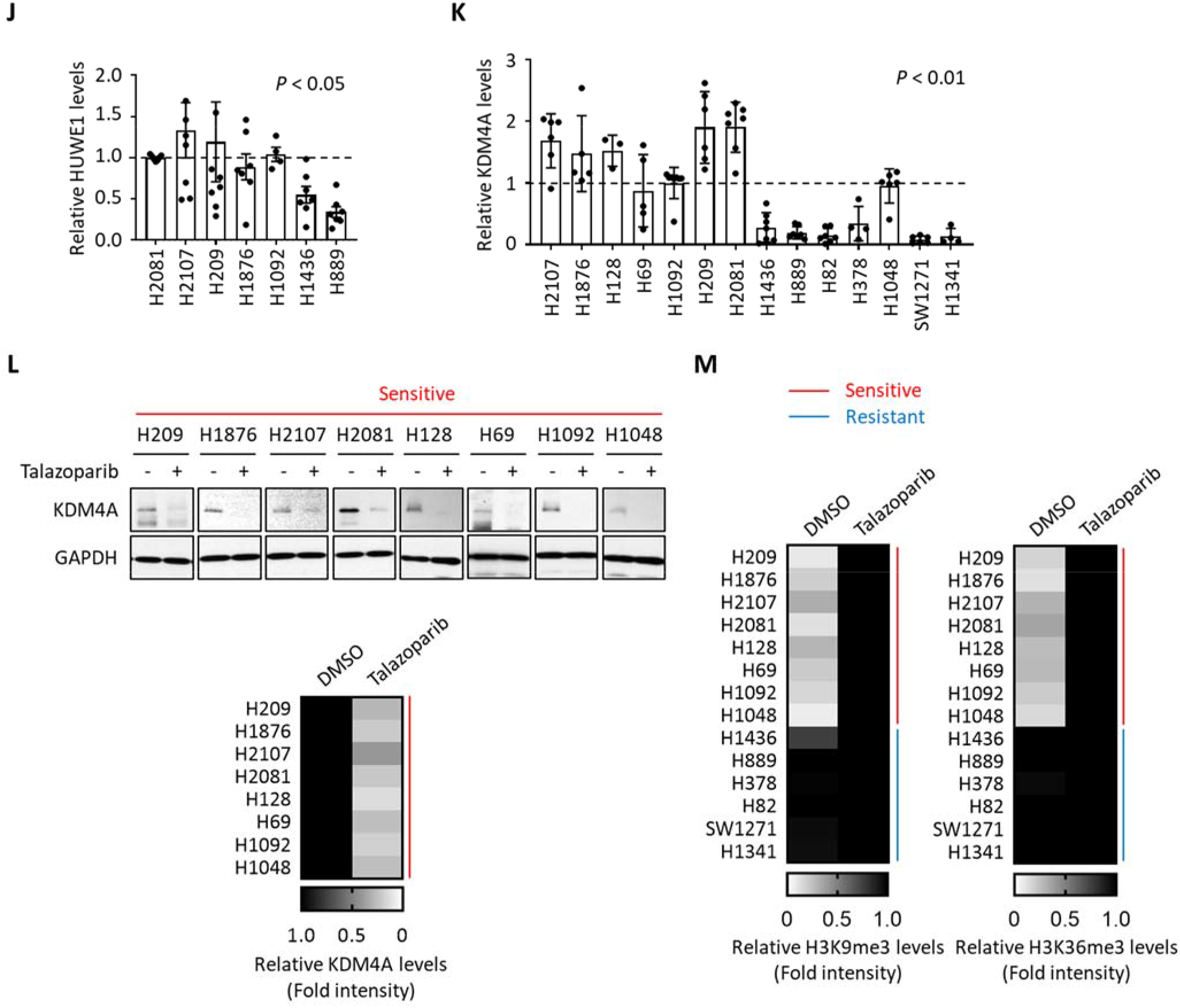
Related to Fig. 6. (A) PARP1 cleavage in a panel of SCLC cell lines treated with talazoparib. A panel of SCLC cell lines were treated with talazoparib (1 (+) and 10 (++) μM for 48 hrs). The cell lysates were subjected to immunoblot analysis and the levels of PARP1 cleavage were normalized. The sensitivity is defined as: Red: sensitive SCLC cell lines; Blue: resistant SCLC cell lines. (B) Viability assays of a panel of SCLC cell lines in talazoparib treatment. A panel of SCLC cell lines were treated with talazoparib (1 (+) and 10 (++) μM for 48 hrs). Viability was measured using a CellTiter-Glo assay and normalized. Red: sensitive SCLC cell lines; Blue: resistant SCLC cell lines. (C) The distribution of SCLC cell lines that are sensitive or resistant to PARPi based on PARP1 cleavage (A) and cell survival ratio (B). Red: PARPi-sensitive SCLC cell lines; Blue: PARPi-resistant SCLC cell lines. (D) The extent of DNA damage in a panel of SCLC cell lines treated with talazoparib. Quantification of the immunoblots presented in Fig. 6A was shown (n = 3). Red: sensitive SCLC cell lines; Blue: resistant SCLC cell lines. (E) The levels of ASCL1, NEUROD1, and POU2F3 in a panel of SCLC cell lines treated with talazoparib. A panel of SCLC cell lines were treated with talazoparib (1 (+) and 10 (++) μM for 48 hrs). The cell lysates were subjected to immunoblot analysis and the levels of ASCL1, NEUROD1, and POU2F3 were normalized. TF^low^, low expression of all three transcription factors. See also Fig. 6B. (F) The levels of ASCL1 and cleaved PARP1 in a panel of ASCL1^high^ SCLC cell lines under a therapeutic concentration of talazoparib. A panel of SCLC cell lines were treated with talazoparib as indicated for 48 hrs and were subjected to immunoblot analysis using the indicated antibodies. (G and H) Heatmap of the normalized levels of ASCL1 (G) and cleaved PARP1 (H) in a therapeutic concentration of talazoparib from Supplementary Fig. 6F. (I) Heatmap of cell survival in a panel of ASCL1^high^ SCLC cell lines under a therapeutic concentration of talazoparib. A panel of ASCL1^high^ SCLC cell lines were treated with talazoparib as indicated for 48 hrs and viability was measured using a CellTiter-Glo assay (n = 3). (J) The levels of HUWE1 in a panel of ASCL1^high^ SCLC cell lines (n = 3-7). Quantification of the immunoblots presented in Fig. 6C was shown. (K) The levels of KDM4A in a panel of SCLC cell lines (n = 3-7). Quantification of the immunoblots presented in Fig. 6F was shown. (L) The levels of KDM4A in a panel of KDM4A^high^ SCLC cell lines treated with talazoparib. A panel of KDM4A^high^ SCLC cell lines were treated with talazoparib (1 μM for 48 hrs). The cell lysates were subjected to immunoblot analysis using the indicated antibodies and the levels were normalized (n = 3). Red: sensitive SCLC cell lines. (M) Histone H3 trimethylation in a panel of SCLC cell lines treated with talazoparib. Quantification of the immunoblots presented in Fig. 6I was shown. Red: sensitive SCLC cell lines; Blue: resistant SCLC cell lines.

**Supplementary Fig. 7.**
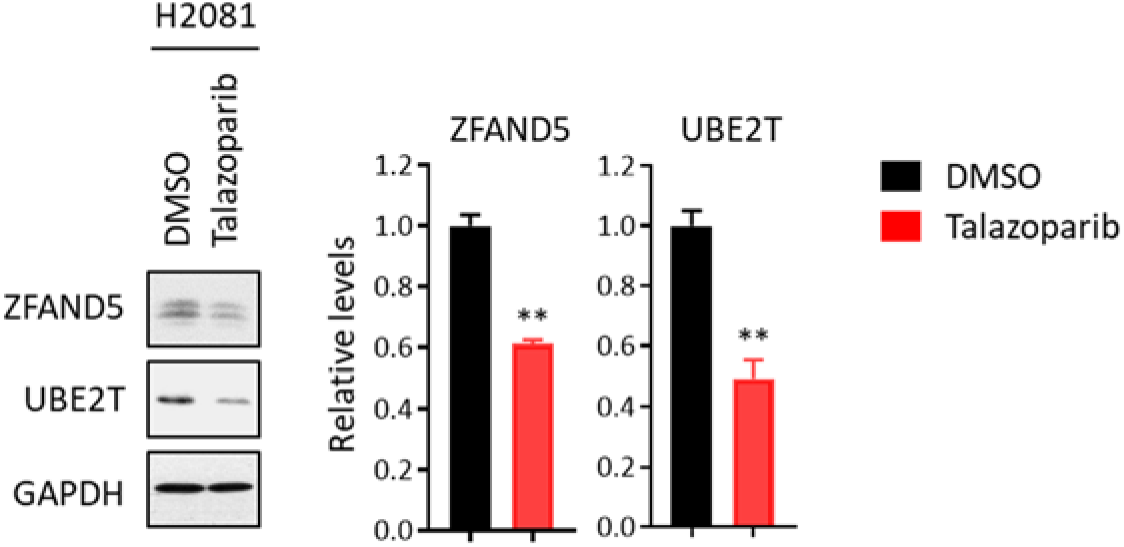
Related to Discussion. (A) The levels of ZFAND5 and UBE2T in talazoparib treatment. H2081 cells were treated with or without talazoparib (1 μM for 48 hrs) and the cell lysates were subjected to immunoblot analysis using the indicated antibodies. Right, the quantification result.

